# A unified theory for organic matter accumulation

**DOI:** 10.1101/2020.09.25.314021

**Authors:** Emily J. Zakem, B. B. Cael, Naomi M. Levine

**Affiliations:** Department of Biological Sciences, University of Southern California, Los Angeles, USA; National Oceanography Centre, Southampton, UK

## Abstract

Organic matter constitutes a key reservoir in global elemental cycles. However, our understanding of the dynamics of organic matter and its accumulation remains incomplete. Seemingly disparate hypotheses have been proposed to explain organic matter accumulation: the slow degradation of intrinsically recalcitrant substrates, the depletion to concentrations that inhibit microbial consumption, and a dependency on the consumption capabilities of nearby microbial populations. Here, using a mechanistic model, we develop a theoretical framework that explains how organic matter predictably accumulates in natural environments due to biochemical, ecological, and environmental factors. The new framework subsumes the previous hypotheses. Changes in the microbial community or the environment can move a class of organic matter from a state of functional recalcitrance to a state of depletion by microbial consumers. The model explains the vertical profile of dissolved organic carbon in the ocean and connects microbial activity at subannual timescales to organic matter turnover at millenial timescales. The threshold behavior of the model implies that organic matter accumulation may respond nonlinearly to changes in temperature and other factors, providing hypotheses for the observed correlations between organic carbon reservoirs and temperature in past earth climates.

## Introduction

Heterotrophic organisms consume organic matter (OM) for both energy and biomass synthesis. Their respiration transforms much of it back into the inorganic nutrients that fuel primary production. Residual OM accumulates into large reservoirs in the ocean, sediments, and soils. Together, these pools store about five times more carbon than the atmosphere and play a central role in global bio-geochemistry^1^. Therefore, the dynamics of OM cycling and accumulation are key to understanding how the carbon cycle changes with climate^1,2,3^.

Standing stocks of OM are comprised of a heterogeneous mix of thousands of compounds, many of which are uncharacterized, with concentrations ranging over several orders of magnitude^4,5,6,7^. Compounds are often conceptually described in terms of a degree of ‘lability’ that correlates with consumption rates, such that labile compounds have low abundances and short residence times in the environment^8,9^. In most biogeochemical models, OM degradation is dictated by simple rate constants rather than explicit consumption by dynamic microbial communities^10,11^. Though significant progress has been made on integrating OM cycling with microbial community dynamics^12,13,14,15,16,17^, we still lack a mechanistic understanding of the ecological controls on OM and its accumulation.

Dissolved OM (DOM) cycling in the ocean has been studied for many decades, making this reservoir ideal for developing a mechanistic framework for OM accumulation. Three hypotheses have been invoked to explain DOM accumulation in the ocean: 1. “*Recalcitrance*”: Compounds may accumulate because they are relatively slowly degraded or resistant to further degradation by microorganisms^8,9,18,19^. This is consistent with observations, theory, and inferences of a wide range of consumption rates and compound ages in the ocean^20,21,22,23,24,25^, as well as in sediments and soils^26,10,11,27,9^. 2. “*Dilution*”: The accumulation may represent the sum of low concentrations of many organic compounds, each having been “diluted” by microbial consumption to a minimum amount^28^. This is supported by evidence that concentrating apparently recalcitrant DOM from the deep ocean fuels microbial growth^29^. Modeling efforts have reconciled observed carbon ages with this mechanism and have interpreted the minimum concentrations as resource subsistence concentrations – the minimum concentrations to which populations can deplete their required resources^17,30^. 3. “*Dependency on ecosystem properties*”: The accumulation may result from a mismatch between OM characteristics and the metabolic capability of the proximal microbial community (e.g. the substrate-specificity of enzymes)^31,32,33,34^. For example, the dispersal of microbial populations, which is controlled by the connectivity of the environment and which may manifest as a stochastic process^35^, can allow for intermittent or sporadic OM consumption events^32,34^. In soils and sediments, some aspects of these hypotheses apply, while other processes also influence the accumulation of OM such as diverse redox conditions and the physical and chemical dynamics of solid organic particles and mineral matrices.

Here, we investigate why OM accumulates using a stochastic model that simulates the complex dynamics of microbial OM consumption. We find that the mechanisms underlying each of the three above hypotheses come into play simultaneously in the model. We develop a quantitative definition of functional recalcitrance that depends on both the microbial community and the environmental context, in addition to substrate characteristics. We demonstrate the model’s ability to explain the accumulation of DOM in the ocean, and because it is grounded in basic principles of microbial ecology, we suggest that this framework can also be extended for application to soil and sediment environments. Finally, the threshold behavior of the recalcitrance indicator suggests nonlinear OM responses to changes in the environment.

### A mechanistic model of organic matter consumption

We develop a model of organic matter consumption by microbial populations using established forms of equations for microbial growth and respiration^36,12^. The model resolves multiple pools of OM (*n* = 1,000) that are supplied stochastically and consumed by one or more microbial populations (*n* = 1,000 or 2,000, Eqns. 4–6, Fig. 1). Stochastic supply captures the variable nature of the release of organic compounds, which is a function of complex biological dynamics (e.g. exudation, lysis, grazing). We represent the net impact of each complex OM-microbe interaction (e.g. hydrolysis, enzymatic rates, cellular allocation of enzyme, free energy released by OM oxidation^37,16,32,33^) with a simplified set of parameters: maximum uptake rate, half-saturation concentration, and biomass yield (Materials and Methods). To include the impact of variable community composition, we modulate the OM consumption by each population over time according to its stochastically assigned probability of presence. We vary the degree of ‘specialists’ (consuming a single OM pool) vs. ‘generalists’ (consuming multiple OM pools), incorporating a penalty that increases with the number of pools consumed to represent a tradeoff among the strategies. We vary both the number of pools consumed by each population and the number of consumers of each pool (Fig. 1, Fig. S1). Population loss rates are proportional to biomass according to both quadratic and linear mortality parameters, simulating predation, viral lysis, senescence, and cell maintenance demand.

**Figure 1:**
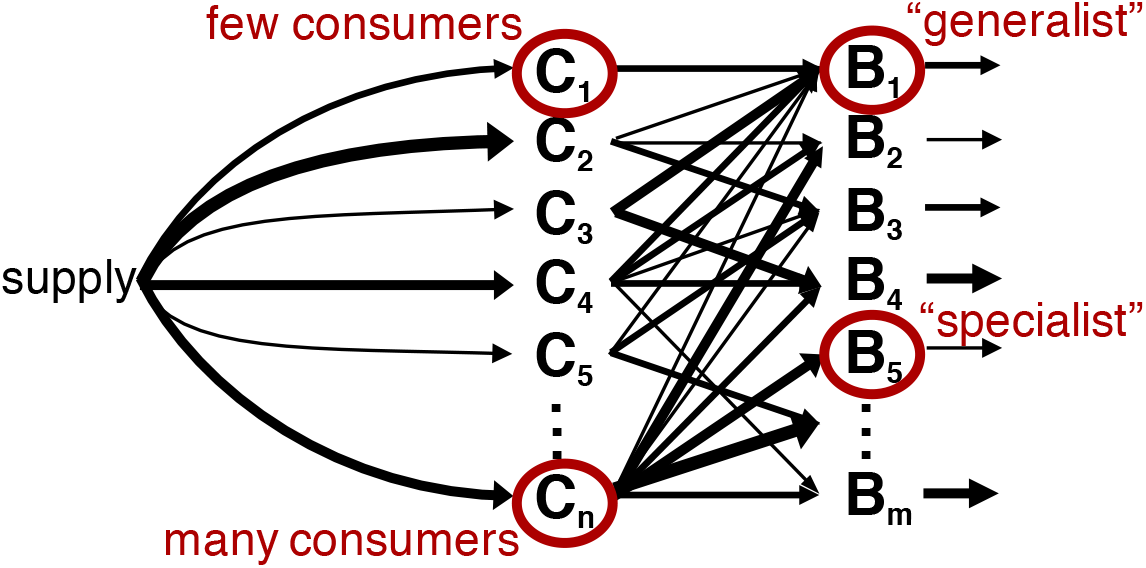
Schematic of the organic matter (OM) consumption model. Multiple OM pools *C* and microbial populations *B* are resolved. The parameter values dictating the supply of each OM pool, the interaction between each pool and the microbial population (uptake kinetics and yield), and the loss of biomass (to viral lysis, grazing, senescence, and cell maintenance) are assigned stochastically. Here we show an illustrative example where the fluxes dictated by these parameter values are represented with different widths of arrows. The supply and the presence or absence of each population vary stochastically over time in the model according to assigned probabilities.

Because we expect the values of these growth and mortality parameters to vary widely among organisms and substrates, we sample all parameter values from uniform distributions over wide, plausible ranges (Table 1, SI Text 1). We numerically integrate the equations forward in time, allowing the concentrations of OM pools to emerge from the ecological interactions. The dynamics presented here are robust across the parameter space, variations in the model structure, and variations in the number of OM pools and populations (SI Text 2 and 3, Figs. S2-S7). Sequential transformation of one OM pool to another due to incomplete oxidation gives qualitatively similar solutions (SI Text 2), although this may increase compound age^17^. We present results from simulations integrated for ten years (Fig. 2).

**Figure 2:**
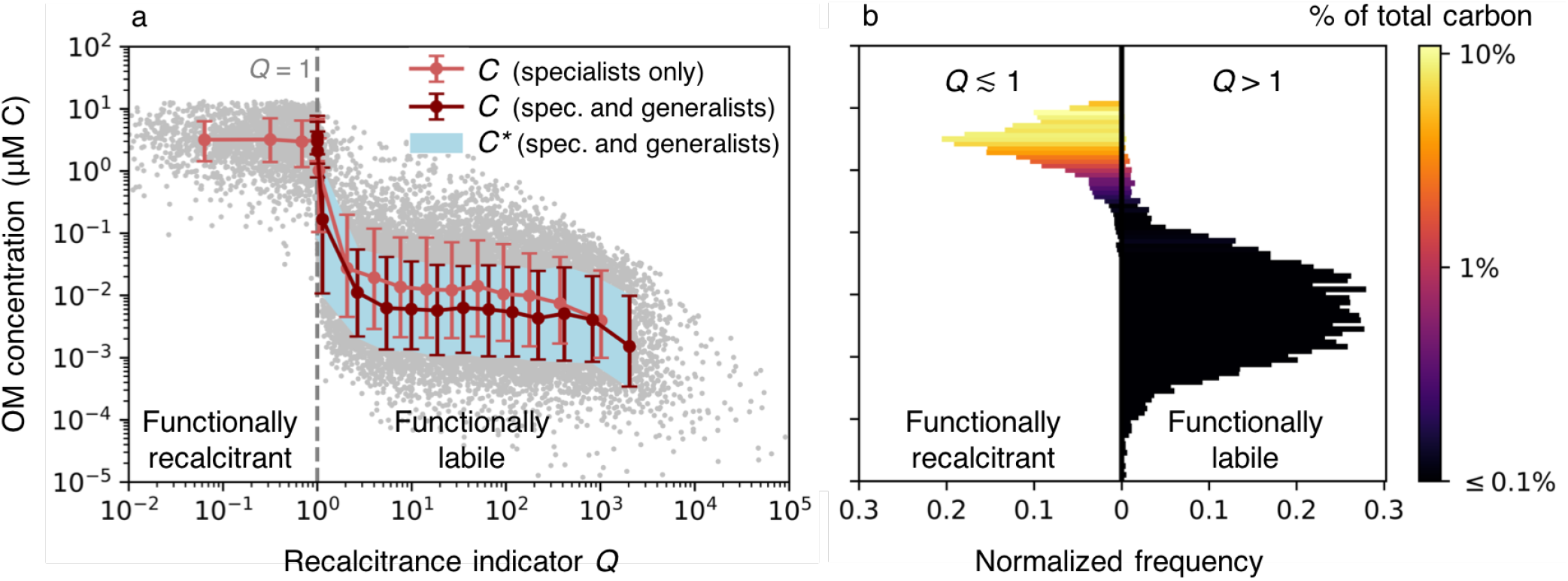
Simulated concentrations from the stochastic organic matter (OM) consumption model. (a) Simulated OM concentrations *C* and associated diagnostic *C*\*, the subsistence concentrations of the microbial consumers (Eqns. 2 and S18), against recalcitrance indicator *Q* (Eqn. 3). The *Q* = 1 threshold (grey dashed line) delineates the functionally recalcitrant (accumulating) and functionally labile (equilibrated) OM. We illustrate compiled results from two model versions, each resolving 1,000 OM pools: one with only 1,000 ‘specialist’ microbial populations, and one with the specialists and an additional 1,000 ‘generalist’ populations, which consume varying numbers of OM pools. We compile ten simulations of each model version so that 10,000 OM concentrations underly the illustrated statistics. The red and light red dots indicate the binned means for the two compilations. The red and light red bars (for the model solutions) and the light blue shaded area (for diagnostic *C*\*) indicate the 16th and 84th percentiles (equivalent to one standard deviation for a Gaussian distribution). The grey dots indicate the 20,000 individual OM concentrations from both compilations combined. (b) Normalized frequencies of the concentrations and their contributions to total carbon in the model (for the version with both specialists and generalists). Frequencies are split at *Q* ≈ 1 (cutoff at 1.01).

**Table 1:** Parameters and their distributions for the OM microbial consumption model. Parameter values are assigned stochastically according to uniform distributions over the indicated ranges.

| Parameter | Symbol | Value (range) | Units |
| --- | --- | --- | --- |
| Number of OM pools* | $n$ | 1,000 | |
| Number of populations<sup>†</sup> | $m$ | 1,000 and 2,000 | |
| Probability of presence | $P$ | 0 – 1 | |
| Total OM supply (all pools) | $\sigma_T$ | 0.1 | $\mu\text{M C d}^{-1}$ |
| Potential supply (each pool) | $\sigma$ | $\sigma_T n^{-1}$ | $\mu\text{M C d}^{-1}$ |
| Probability of supply | $q$ | 0 – 1 | |
| Maximum specific uptake rate | $\rho^{\max}$ | $10^{-2} - 10^{2\ddagger}$ | $\text{d}^{-1}$ |
| Half-saturation concentration | $k$ | $\rho^{\max}(10^0 - 10^2)^{-1\ddagger}$ | $\mu\text{M C}$ |
| Yield (growth efficiency) | $y$ | 0 – 0.5 | $\text{mol C mol C}^{-1}$ |
| Quadratic mortality rate | $m^q$ | 0.1 – 1 | $(\mu\text{M C d})^{-1}$ |
| Linear mortality rate | $m^l$ | 0 – 0.01 | $\text{d}^{-1}$ |
| Loss rate | $L$ | $m^q B + m^l$ | $\text{d}^{-1}$ |

\*Here, we illustrate two 10-member ensembles with 1,000 OM pools in each individual model, giving a total of $10^4$ pools. See Fig. S3 for a model with $10^4$ pools.
<sup>†</sup> The community consumption matrix dictates which populations consume each pool (Fig. S1). See Figs. S4–S7 for variations, including variations in the ratio of populations to pools from 2:1 to 1:1,000.
<sup>‡</sup> Varied over a log rather than a linear range.

The solutions reveal a bimodal distribution of OM concentrations (Fig. 2), implying a set of qualitatively distinct controls on OM accumulation. Whether or not the bimodality is discernible depends on the parameter distributions (Fig. S13), as well as other sources and sinks not included in the model (e.g. photolysis). In the simple model, the majority of pools are depleted to relatively low concentrations (10^-4^ to 1 μM C), while a subset accumulates to substantially higher concentrations (0.1 to 10 μM C). The latter accumulated pools comprise the bulk of total carbon content (Fig. 2b).

### Diagnosing functional recalcitrance

We evaluate whether each OM pool equilibrates or accumulates in the model. This indicates whether the pool can sustain a microbial population in the given environment, and we classify the pool as functionally labile or recalcitrant accordingly. For example, we describe the population dynamics of specialist population *j*, subsisting solely on OM pool *i*, as (from Eqns. 4 and 6):

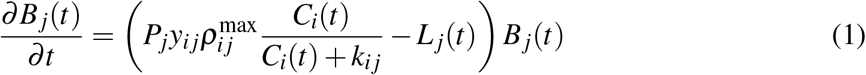

where *B_j_* is the biomass, *P_j_* is the probability of the presence of population *j*, *y_ij_* is the biomass yield, 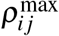 is the maximum uptake rate, *k_ij_* is the half-saturation concentration for uptake, *C_i_* is the concentration of the OM pool, and *L_j_* is the population loss rate, which varies as a function of the biomass (Eqn. 6, Table 1). When the system is at or close to steady state 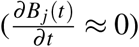, the concentration of pool *i* can be estimated as:

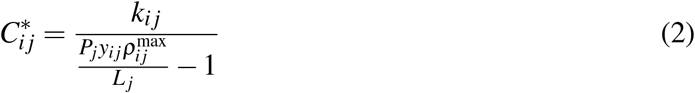

which is the subsistence concentration of OM pool *i* for specialist population *j*^30^. For a pool with multiple competing consumers, the concentration of that pool will be set by the population with the lowest subsistence concentration for that pool^30^. The population can then continue to consume the pool in proportion to its supply while maintaining the subsistence concentration^30,17^.

For the OM pool to equilibrate (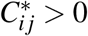 in Eqn. 2), the maximum rate of local biomass synthesis 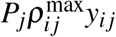 must exceed *L_j_*, which is the population turnover rate at steady state. Using 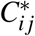 as a diagnostic, and extending the expression to generalists that can consume more than one OM pool (Eqns. S17 and S18), we find that many concentrations of the modeled pools precisely match the minimum subsistence concentration among their consumers, and thus have equilibrated (Figs. 2a, S8 and S9). Because these pools sustain microbial growth in this particular model environment, we term these pools ‘functionally labile’. These low concentrations are consistent with the measured nanomolar or lower concentrations of known labile constituents of marine dissolved organic matter, such as free amino acids and glucose^4,38^.

Most of the pools that accumulate to higher concentrations never equilibrate in the simple model. For these pools, the loss rates of all consuming populations match or exceed their maximum biomass synthesis rates. As a result, no populations can be sustained on these pools alone in this model environment. We term these pools ‘functionally recalcitrant.’ We can robustly define the threshold where pools transition from being functionally labile (depleted to 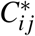) to functionally recalcitrant (accumulating). We define a recalcitrance indicator *Q_i_* for pool *i* as:

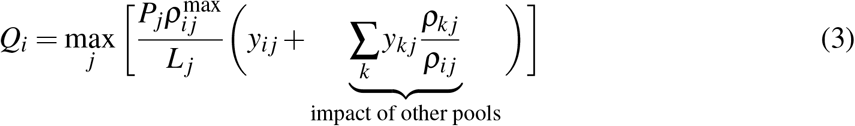

where index *k* denotes a pool other than pool *i* consumed by generalist population *j*, and *ρ_kj_*/*ρ_ij_* is the relative uptake of pool *k* to pool *i* by population *j* (see SI Text 4 for derivation). For specialists, the term representing the impact of other pools drops out of the equation. If *Q_i_ >* 1, pool *i* is functionally labile: at least one population can deplete it to its subsistence concentration given sufficient time, with the equilibration timescale dictated by the associated growth and mortality parameters. If *Q_i_* ≤ 1, pool *i* is functionally recalcitrant, and it accumulates over time in our model. Thus *Q_i_* = 1 serves as an emergent threshold between functional lability and functional recalcitrance (Fig. 2).

The recalcitrance indicator *Q_i_* demonstrates how recalcitrance is simultaneously governed by chemical, biological, ecological, and environmental characteristics. In Eqn. 3, an enzyme-dependent substrate-microbe interaction *i-j* is captured by both *y_ij_* and 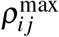, which also reflect the energetic content and the accessibility of the OM^39,40,41^. The encounter probability of the population with the OM pool (*P_j_*) and the biomass loss rate (*L_j_*) capture the ecological context - the diversity and abundances of the local microbial populations, predators, and viruses. Many factors control these processes, including selection and environmental connectivity^35^, which is shaped in part by physical conditions such as circulation and sinking particles in the ocean and porosity and diffusion in soils. Diversity and connectivity also modulate the availability of other pools for uptake by generalists. In Eqn. 3, the uptake of an additional OM pool *k* by population *j* can increase its potential to deplete pool *i* (i.e. *Q_i_* increases). In other words, the ability of consumers of OM pool *i* to consume other pools increases the functional lability of pool *i*. This provides a mechanistic explanation for the observed ‘priming effect’ in which the addition of other substrates allows for the metabolization of a given pool^42,43,34^.

In the environment, a functionally recalcitrant OM pool may accumulate or diminish at a rate dependent on production, consumption, and physical transport over time, or it can equilibrate due to an abiotic, concentration-dependent sink such as photolysis^8,34^. In the model version with many generalists, *Q_i_* reaches a minimum of one (to within 1%) (Fig. 2a). When *Q_i_* ≈ 1, pools are unequilibrated and functionally recalcitrant, but consumption can continue by consumers whose loss rates have dynamically adjusted to approach their maximum biosynthesis rates.

Recalcitrance emerges as a community- and context-specific phenomenon that can change in time and space (Fig. S10). Critically, the recalcitrance indicator for each OM pool (*Q_i_*) is defined as the maximum of multiple population-specific values (Eqn. 3) - one for each population *j* that consumes pool *i*. Consequently, whether each pool is functionally labile or recalcitrant depends on the local microbial community. For a diverse community of consumers, we can analyze the fraction of the community that experiences each pool as recalcitrant (Fig. S11). This community dependency implies that statistically, functional recalcitrance may be more prominent when OM is exposed to a lower diversity of heterotrophic microorganisms. This also implies that functional recalcitrance may arise from the requirement for specialized enzymes or expensive consumption pathways for some types of OM^44^: if specialization is required, there may be less possible consumers overall, and so it becomes less likely that any one consumer is present in the given environment. This is consistent with evidence that specific heterotrophic clades consume carboxyl-rich alicyclic molecules (CRAM), which comprise a significant fraction (up to 8%) of marine DOC^6,45^.

### Unification of hypotheses

The three current hypotheses for dissolved OM accumulation in the ocean – recalcitrance, dilution, and dependency on ecosystem properties – each explain aspects of the total amount of carbon in the model. Additional processes, such as mineral protection and diverse redox conditions of soils and sediments, can also be incorporated into the framework to modify it for these other systems. We may consider each hypothesis individually as a limit case for the formation of large organic carbon reservoirs in natural environments. Total organic carbon content is the sum of all OM pools. A traditional view of recalcitrance, focused on intrinsic qualities of the substrate or of the microbe-substrate-specific reaction, is represented in the model by the biomass yield *y_ij_* and maximum uptake rate 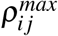. The quality of electron acceptor or mineral protection can also be represented by these parameters. As 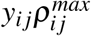 becomes small, all else the same, OM becomes recalcitrant (Eqn. 3) and the total organic carbon pool becomes large. The number of OM pools *n* that are present can impact total carbon in two opposing ways. As *n* increases, total carbon increases even for low, equilibrated subsistence concentrations (the “dilution” hypothesis). However, the impact of other OM pools (priming) means that as *n* increases, the likelihood of *Q_i_* > 1 increases, decreasing the likelihood of functional recalcitrance and thus potentially decreasing total carbon. Dependency on ecosystem properties is encapsulated in the population turnover rate *L_j_* and the probability *P_j_* of co-occurrence with consuming populations. As *L_j_* increases, OM becomes recalcitrant and total carbon increases. As the number of nearby consumers increases, the likelihood of *Q_i_* > 1 increases. Thus as *P_j_* becomes small, total carbon becomes large.

The degree to which each mechanism controls OM accumulation in different environments therefore depends on the parameter space that sets the population and OM characteristics. Here, we assume uniform distributions for these parameters using plausible ranges for the ocean (Table 1, SI Text 1). These ranges will vary with the environment. For example, if stochasticity in population presence does not apply to a given sediment ecosystem, then probability of presence *P* may be set to one for analysis of that environment. The model is consistent with that of ref.^17^ in that intrinsic recalcitrance is not necessary for OM accumulation in the ocean, as well as with experimental evidence for dilution-limited consumption (SI Text 5, Fig. S12)^29^. Here we provide a generalized framework that encapsulates a more complete set of dynamics than in ref.^17^ - one that is also consistent with evidence of recalcitrance^8,18,19^ and the impact of the microbial community^31,33,44,45,34^. The emergent distributions of OM degradation rates are consistent with theory and observations that remineralization rates are lognormally distributed over a wide range due to multiplicative stochasticity in the underlying processes^27^ (Fig. S13). They are also consistent with continuum intrinsic reactivity models^11,10^ for which the assumed rate distributions tend to be similar to lognormal over the bulk of the distributions^46^.

### Predicting OM accumulation patterns

Our framework can help explain large-scale patterns in OM accumulation. Here, we use our model to understand the vertical structure of dissolved organic carbon (DOC) in the ocean. Globally, DOC concentrations peak at the sea surface and approach a minimum at depth (Fig. 3a)^8^. Since the stochastic model is not practical for multi-dimensional biogeochemical models, we utilize a reduced-complexity model analog that captures the essence of the stochastic model, resolving 25 aggregate pools. We incorporate this model analog into a fully dynamic ecosystem model of a stratified marine water column, where production and consumption of all organic and inorganic pools are resolved mechanistically as the growth, respiration, and mortality of photoautotrophic and heterotrophic microbial populations (SI Text 6, Fig. S14). The model is integrated for 6,000 years to quasi-equilibrium (SI Text 6).

**Figure 3:**
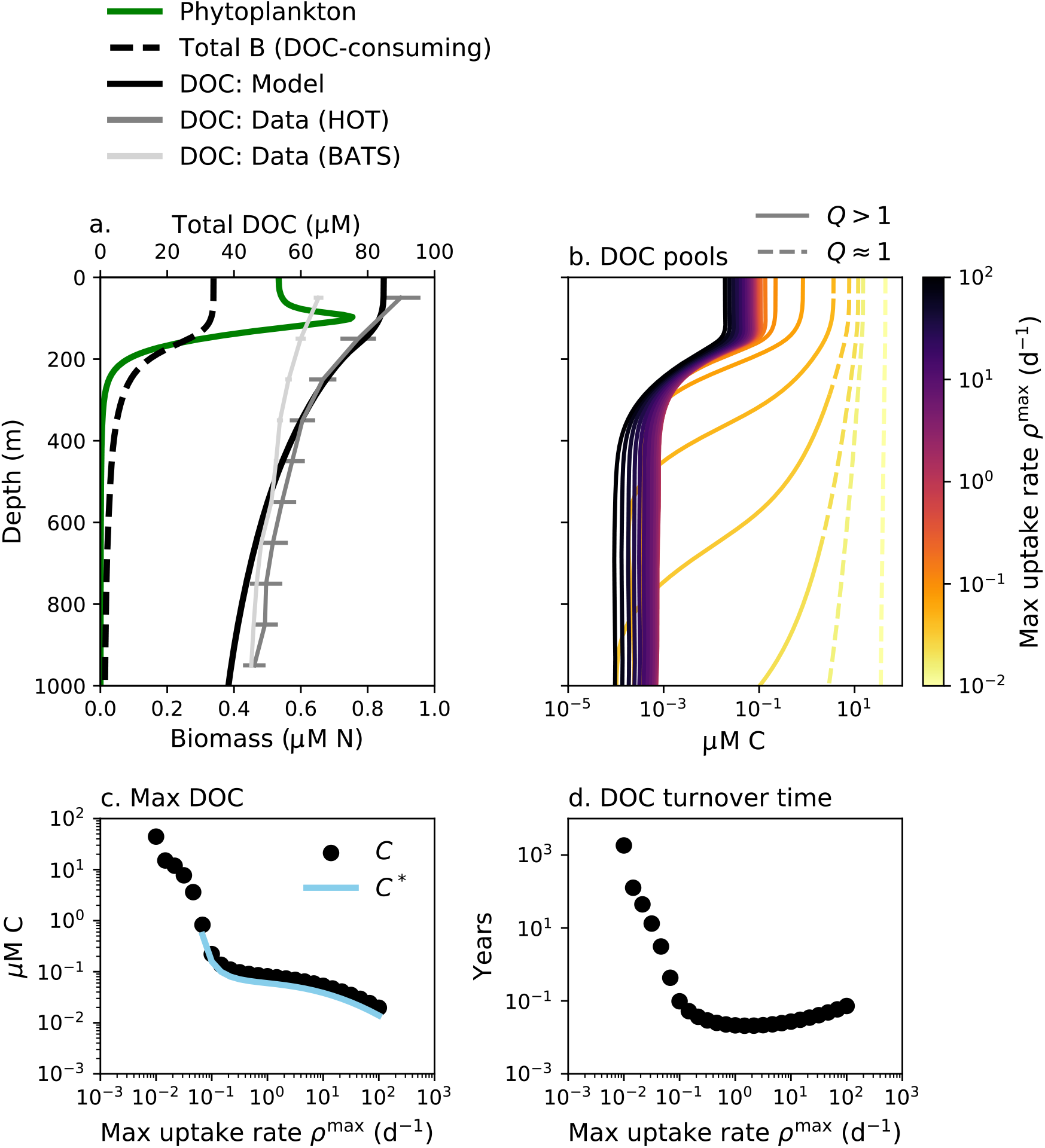
Solutions for the marine ecosystem water column model, which simulates the accumulation of dissolved organic carbon (DOC) in the ocean as a function of depth. (a) Phytoplankton biomass, total DOC-consuming biomass B, and total DOC. Annual average profiles of total DOC from two open ocean time series stations are illustrated: HOT (the Hawaii Ocean Time-series in the Pacific Ocean) and BATS (the Bermuda Atlantic Time-series Station in the Atlantic Ocean)^49^. (b) The concentration of each of the 25 resolved DOC pools, which are differentiated in the water column model by maximum uptake rate *ρ*^max^ (color scale). Each pool is categorized as functionally labile (solid line) or functionally recalcitrant (dashed line) as a function of depth using recalcitrance indicator *Q* (Eqn. 3). (c) The maximum (surface) concentration *C* of each DOC pool and the associated diagnostic *C*\*, the subsistence concentration of the microbial consumer population (Eqn. 2), plotted against the maximum uptake rate for that pool. (d) The turnover time of each DOC pool calculated diagnostically as the quotient of the integrated concentration and the integrated consumption rate, plotted against maximum uptake rate.

Ecological interactions in the model result in characteristics typical of a marine water column (Fig. 3, Fig. S15). Modeled DOC accumulates throughout the water column. Total DOC decreases smoothly with depth, with higher surface DOC transported to depth by vertical mixing (Fig. 3a). Most pools are depleted to subsistence concentrations throughout the water column (Fig. 3b,c). One pool remains functionally recalcitrant throughout the entire water column due to its slow consumption rate (lightest yellow line in Fig. 3b), which is consistent with the observed homogenous composition of aged marine DOC ^47,7^.

Many DOC pools in the model accumulate at the surface and become depleted at depths of 500 to 1,000 m. This transition is due to the increase in *Q* from the surface to depth (Fig. 3b). Productivity peaks in the sunlit surface, and so total biomasses, activity rates, and therefore population loss rates reach maximum values at the surface and attenuate with depth. This transition is consistent with observations that a subset of DOC is resistant to consumption by surface communities but able to be remineralized by deep communities^31^. Thus an ecologically determined transition from functional recalcitrance to functional lability, dictated by the sustainability of microbial populations represented in aggregated form, explains much of the decrease in DOC with depth. The subsistence concentrations also decrease with depth as loss rates decrease, and thus functionally labile OM also contributes to the vertical DOC gradient.

Our framework may also be employed to investigate microbial control on OM in soils and sediments. The model can be adapted to incorporate the different characteristics of these environments. For example, we here employ a simple parameterization for the supply of each OM class, but a sediment or soil model version could include more sophisticated descriptions of how the physics and chemistry of solid particles and mineral matrices impact the supply rate. Though Michaelis-Menten uptake kinetics do not apply to the enzymatically catalyzed degradation of polymer to monomer form, the ecological principles of our framework should still hold (SI Text 7). Indeed, we find that even in its current form, the simple model captures a key observation of sediment OM: the proportional increase in OM decomposition rate with increased OM concentration^48^ (Fig. S16). This further demonstrates consistency with the predictions of established first-order kinetic decomposition models^48,12^. Future work can also explore the impact of more enzymatically diverse sediment pelagic microbial communities ^33^ by experimentally altering the community consumption matrix to include a greater degree of ‘generalist’ ability. Also, the model may incorporate diverse redox conditions with a yield or uptake rate that varies with the electron acceptor. This may suggest that some types of OM are functionally labile when oxygen is available, but functionally recalcitrant in anoxic environments.

### Implications

Our model is consistent with the observations that the majority of the diverse types of OM is present at relatively low (< 1 μM) concentrations, while the majority of the total standing stock is functionally recalcitrant^8^ (Fig. 2b, Fig. 3b). This is also consistent with the conclusions of previous continuum reactivity models^10,50^. Recalcitrant OM pools may equilibrate if subjected to abiotic concentration-dependent sinks^8^, or they may change slowly with time^7,8^. Our framework further emphasizes that apparently slow consumption rates of recalcitrant DOC in the ocean may be controlled by the frequency of encounter of ‘the right’ populations and substrates, in addition to cellular processing limitations. This is consistent with the understanding that localized sinks cause the 10% to 20% decrease in deep ocean DOC along the deep ocean circulation pathway^32^.

The water column model links the millenial timescales of OM turnover^24^ to microbial consumption occurring on subannual timescales (Fig. 3d). Although explicitly tracking organic transformation through a complex interaction network can also explain old carbon ages^17^, slow change is consistent with inferences that the size of organic carbon reservoirs does not reach a steady state over geologic timescales^2^. While our model is compatible with the “dilution” hypothesis, it also incorporates these other explanations, and so is consistent with a broader set of observations, including the compositional uniformity of ubiquitous recalcitrant classes^47,7^.

A key aspect of our framework is the threshold behavior of the accumulation. The threshold, *Q* = 1, is set by the dynamics of the microbial populations that consume the OM pools. *Q* = 1 represents an ecological threshold along a continuum of OM and microbial characteristics that encompasses a variety of mechanistic factors, including those known to influence recalcitrance such as thermodynamic limitations^39^, enzymatic control^33^, mineral protection^40,43,51^, and molecular properties^19^. The nonlinear behavior of the threshold suggests that small changes in the environment can drive large depletions or accumulations of OM.

Consumption of recalcitrant OM depends on the rate of microbial processing, which increases with temperature. All else the same, the model predicts that less OM accumulates at higher temperatures (SI Text 3, Fig. S10c). Indeed, the loss of soil OM is a likely positive feedback to current warming^52^. The framework here additionally suggests that a decrease in OM with warming may be nonlinear due to some of the pools crossing the threshold from functionally recalcitrant to functionally labile (Fig. S10c). This may help to understand the correlations between temperature and organic carbon reservoirs in past earth climates, such as increased ocean carbon burial, “inert” soil carbon reservoirs, and perhaps marine DOC during glacial periods^53,54^. Temperature-driven nonlinearity may also constitute an explanation for the ten-fold higher microbial utilization rates of DOC in the warmer deep Mediterranean compared to the colder deep open ocean^55^. Using this framework to quantitatively predict changes in organic carbon reservoirs with current increases in global temperature will require accurate estimates of microbial community loss rates, as well as an understanding of how temperature will impact both these microbial rates and the diversity of the community.

We identify a set of controls on OM accumulation and turnover rooted in the complexity of microbial ecosystems. Previously disconnected hypotheses for OM accumulation, including the many mechanisms giving rise to functional recalcitrance, are subsumed within one framework. OM concentrations are mediated by the characteristics of substrate-microbe interactions, the heterogeneity of organic substrates, microbial community dynamics, and the ecological and biogeochemical diversity set by the connectivity of the environment (Fig. 4). The model is consistent with a comprehensive set of observations and theory of OM concentrations, turnover rates, and ages. The framework can be used to quantify the degree to which each of the subsumed hypotheses explains OM accumulation in different environments and to develop testable hypotheses for how organic reservoirs change with the biogeochemical environment.

**Figure 4:**
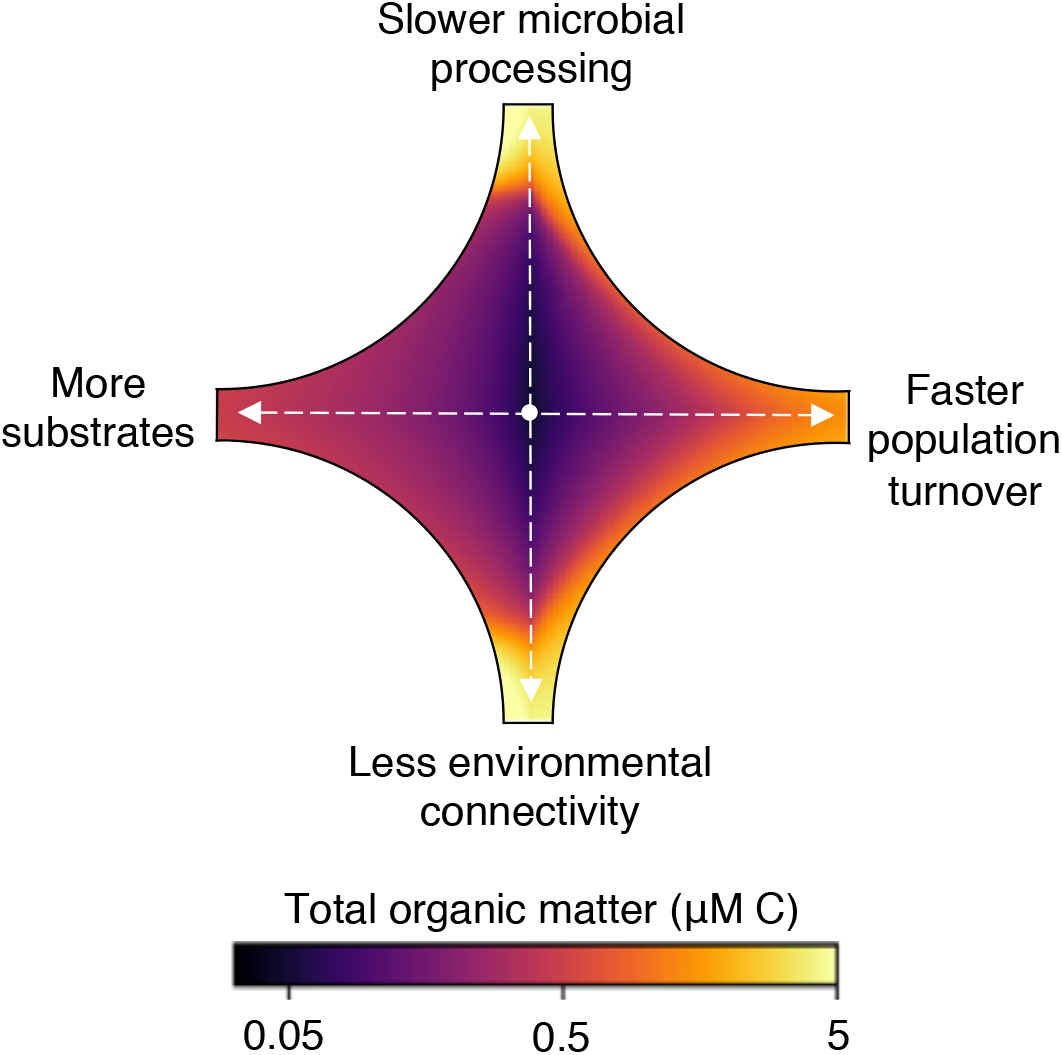
Summary of controls on organic matter accumulation by microbial consumption. Starting from a representative, arbitrary concentration in the center, the change in total organic matter carbon is calculated for a ten-fold change in each of four parameters (i.e. two parameters vary in each quadrant): slower microbial processing via a reduced maximum uptake rate, faster turnover via an increased population loss rate, less connectivity via a reduced likelihood of population presence, and more substrates (chemical diversity) via a greater number of OM pools.

## Methods

### Model equations

We describe microbial consumption and growth on pools of organic carbon. The model framework is sufficiently general to also account for inorganic nutrients, and may be extended to account for the cycling of other elements. We model the uptake *ρ_ij_* of each OM pool *i*, according to its concentration *C_i_*, by microbial population *j* as a function of time *t* using a saturating (Michaelis-Menten) form as:

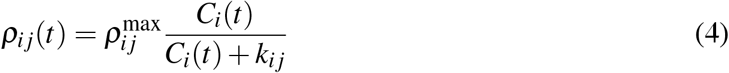

where 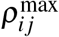 is the maximum uptake rate and *k_ij_* is the half-saturation constant (Table 1).

Each population synthesizes biomass according to a growth efficiency for each pool (yield *y_ij_*), and loses biomass at a rate proportional to its biomass according to a quadratic mortality parameter 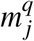 (implicitly representing higher order predators and viruses) and linear mortality parameter 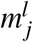 (representing cell maintenance demand and senescence). The rates of change of the concentration *C_i_* of pool *i* and the biomass *B_j_* of population *j* are:

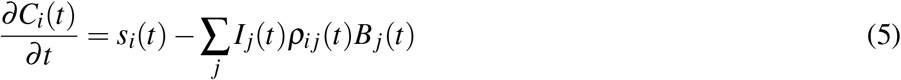

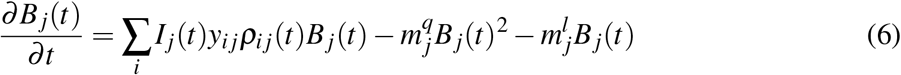

where *s_i_*(*t*) is the supply rate of pool *i*, which is governed by the probability of the supply of each) pool *q_i_* as

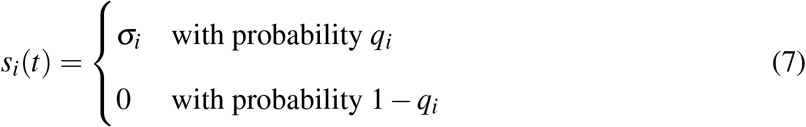

for potential supply rate *σ*_i_, which here is a fraction of total OM supply to the domain (Table 1). The term *I_j_* (*t*) indicates the presence of population *j* at time *t* according to the probability of presence *P_j_* (see detail below) as

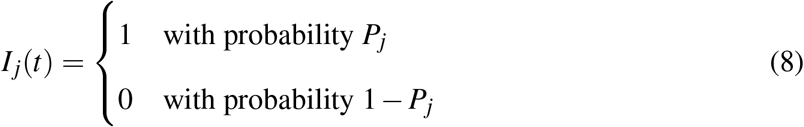

Because the presence of population *j* averages to *P_j_* over time, we include *P_j_* in Eqns. 1-3 for conciseness. All parameter values (*ρ*^max^, *k* (via the affinity *ρ*^max^k^-1^), *y, m^q^, m^l^, q*, and *P*) are set by randomly sampling from uniform distributions (Table 1, SI Text 1).

Yield *y_ij_* reflects the cost of each enzyme and the free energy released by OM oxidation. 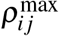 and *y_ij_* may be interdependent due to cellular optimization strategies, reflecting inherent tradeoffs between protein allocation and efficiency^56,57^. The varying combinations of 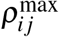 and *y_ij_* can also represent the different modes of uptake of high molecular weight dissolved OM^58^. Analogously, the different parameter combinations can account for the additional feedback between the external concentration and the rate of cellular processing 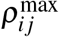. Real populations may change their cellular machinery due to plasticity, where, in the model, the many sets of parameters represent the static phenotypes among these different modes.

### Probability of presence

Observations show that community composition dictates the character of DOM remineralization in seemingly unpredictable ways^60,61^, and evidence supports localized sinks of deep DOM^32^. To simulate this impact, we include a population presence-absence dynamic in the model. Each population is assigned an overall probability of presence *P_j_*, simulating the sporadic presence of rare functional types and the nearly guaranteed presence of ubiquitous types. When *I_j_* (*t*) = 0, the population does not consume OM or synthesize biomass at that timestep, but its biomass is still subject to loss. In effect, this dynamic extends the range of maximum processing (synthesis) rates to lower values, demonstrating how the absences of particular functional types contribute to longer effective remineralization timescales. This dynamic results in the majority of interactions at intermediate (though still widely ranging) rates, with very slow and very fast interactions being fairly rare (Fig. S13). Over time, the average presence approaches *P_j_*, and the steady state balances calculated with the overall probability *P_j_* closely match the model solutions.

### Consumption matrix

A consumption matrix dictates which populations consume which OM pools (Fig. S1). We vary the ‘specialist’ vs. ‘generalist’ capabilities of the populations with respect to the number of OM pools taken up by each population (*n_up_*, which can vary from one to n, the number of OM pools), as well as with the widespread popularity of each pool with respect to the number of consumers of each (*n_cons_*, which can vary from one to *m*, the number of populations) (Fig. 1). In the model version with solely specialists (Fig. 2, Fig. S2), each population consumes only one unique pool. For the generalist populations (Fig. 2, Fig. S2), we first randomly assign *n_up_* to each population drawing from the linear range from one to *n*. Second, we assign a weight to each OM pool of its probability of being consumed *n_cons_*, varying the weights linearly. Finally, we assign the specific pools taken up by each population (i.e. we fill each column of the consumption matrix) by sampling from the *n* possibilities with the weights. For the weighted sampling, we use the algorithms “ProbabilityWeights” and “sample” in the StatsBase package in Julia.

### Simulations

In the simulations illustrated in Fig. 2, we resolve 1,000 OM classes and 1,000 or 2,000 pools of biomass: a model version with 1,000 specialists, and a model version with the 1,000 specialists and an additional 1,000 with a range of generalist ability. Results with the latter 2,000 pools are similar to a model with only the 1,000 generalists. For each experiment, we run the model forward in time for ten years, until the pools that have the potential to equilibrate have equilibrated. The concentrations of many of the recalcitrant pools continue to increase over time (unless a concentration-dependent sink is added to the model). Fig. S13 illustrates the resulting distributions of biomass concentrations, OM concentrations, and remineralization rates of the ensembles, which are consistent with observed and inferred distributions of OM characteristics, remineralization rates, and ages in the ocean, sediments, soils, and lakes^21,26,62,10,27,63,64,22,23,24,25^.

In SI Text 2 and Figs. S2-S7, we demonstrate the qualitative consistency of the solutions across variations of the model. All simulations support the conclusions presented in the main text. Solutions vary quantitatively but not qualitatively with variations in the generalist capabilities of the microbial populations, the number of OM pools resolved, the ratios of OM pools to populations resolved, the time of numerical integration, and the mode of uptake by the populations (additive consumption vs. switching over time to optimize growth). In the model version where generalists switch their consumption over time to optimize synthesis (Fig. S7), values of *Q* < 1 result for some pools as generalists cease to consume functionally recalcitrant pools despite their capability to do so.

### Reduced-complexity model version

For the reduced-complexity model of OM consumption used in the marine ecosystem model, we collapse the complexity onto one master ‘lability’ parameter – the maximum uptake rate – and we resolve fewer OM pools (*n* = 25) (Fig. S15). The values of *y, m_q_, m_l_*, and uptake affinity are kept constant, since their variation affects the solutions quantitatively but not qualitatively. A specialist population, which represents multiple clades in aggregate, consumes each pool. Since we don’t include stochastic processes, the probability of presence *P* = 1 for all populations. In accordance with theory and our stochastic model results^65,27^, we assume a lognormal distribution for the partitioning of total OM production into the 25 pools (Fig. S15d), which represents the average outcome of microbial transformation over time and space.

### Marine ecosystem model

The reduced-complexity model version is incorporated into a dynamic marine ecosystem model of a stratified vertical water column, where the production and consumption of all organic and inorganic pools are due to the growth, respiration, excretion, and mortality of microbial populations (Fig. S14). Light and vertical mixing attenuate with depth. Two populations of phytoplankton convert dissolved inorganic carbon and nitrogen into biomass using light energy. Populations of microbial heterotrophs consume DOM (25 pools) and POM (one pool), oxidize a portion of the carbon for energy, and excrete inorganic carbon and nitrogen as waste products. For simplicity, POM is resolved as one aggregate pool sinking at a constant rate. Total DOM is produced from POM degradation (due to the extracellular hydrolysis of POM) and the biomass loss of all populations. The model is a modified version of a published model in which carbon and nitrogen of the organic pools and the biomasses are each resolved independently^66^. Parameter values are listed in Table S1. See SI Text 6 for model equations and further detail.

## Acknowledgments

We acknowledge funding from the Simons Foundation: The Simons Collaboration on Principles of Microbial Ecology (PriME #542389 to N. M. L.) and the Simons Postdoctoral Fellowship in Marine Microbial Ecology (to E. J. Z.), the National Environmental Research Council (NE-R015953-1 to B. B. C.), and the European Union Horizon 2020 Research and Innovation Programme (grant #820989 to B. B. C.). We thank R. Letscher for providing the DOC data compilations and N. Norris, J. McNichol, E. McParland, and the Levine group for comments on the manuscript. The work reflects only the view of the authors; the European Commission and their executive agency are not responsible for any use that may be made of the information the work contains.

## Notes

### Competing Interest Statement

The authors have declared no competing interest.

