## Supporting Information for "A unified theory for organic matter accumulation"

Contents:

SI Text 1: Parameter distributions
SI Text 2: Variations in model structure
SI Text 3: Sensitivity to parameter values
SI Text 4: Steady state balances
SI Text 5: Simulation of Arietta experiments
SI Text 6: Marine ecosystem model detail
SI Text 7: Considering the particle to monomer transition

### SI Text 1: Parameter distributions

We assign parameter values in the model by drawing from wide ranges using the random number generator "rand" in Julia. These distributions are informed by estimating plausible ranges in real microbial communities as listed in Table 1 and described below. Different distributions of parameter values result in different distributions of equilibrated vs. recalcitrant OM pools (Fig. S13). We emphasize that the framework provided by $Q$ is applicable to alternative distributions, and thus that the model solutions remain qualitatively robust across different parameter spaces.

We vary the maximum uptake rate $\rho^{\max}$ over four orders of magnitude, drawing from $10^{-2}$ to $10^{2}$ d$^{-1}$ (Table 1). $\rho^{\max}$ is meant to encompass not just enzymatic uptake rates, but also the net impact of other processes involved in the consumption of organic substrates, such as hydrolysis as well as internal processing constraints. $\rho^{\max}$ sets the growth rate ($y\rho^{\max}$ is the maximum biomass synthesis, or growth, rate), and so its variation represents the wide range of observed microbial growth rates and population turnover rates[1,2]. Specifically, a recent study estimates the average growth rate of marine heterotrophic bacteria of about $0.1 \pm 0.1$ d$^{-1}$[1], while noting that growth rates of some organisms can exceed 100 d$^{-1}$. The lower limit of $10^{-2}$ d$^{-1}$ likely does not capture the lowest rates in sediments – evidence suggests that sediment microorganisms turnover at rates of years to hundreds of years[2] – though mortality rates there are also likely significantly lower.

We vary the uptake affinity $\rho^{\max}k^{-1}$ by two orders of magnitude (Table 1), resulting in variation of half-saturation concentration $k$ of about six orders of magnitude, from $10^{-4}$ to $10^{2}$ µM C. This is consistent with observed ranges of the uptake affinity and half-saturation constants of phytoplankton and heterotrophic microorganisms[3,4,5].

The yield (or the growth efficiency) and the mortality constants are each sampled from linear ranges (Table 1). The upper limit to the yield is 0.5 to reflect the generally lower ranges of net heterotrophic carbon utilization efficiencies in natural environments compared to the laboratory[6,7]. The mortality constants are largely uncertain, but model results remain qualitatively robust for different values of these parameters. The quadratic mortality parameter largely influences the resulting concentrations of sustainable biomass, impacting the population loss rate. The linear mortality rate constitutes the lower limit to a population's sustainability in its environment, representing the cell's maintenance demand. Thus the maintenance demand may be lower in environments such as sediments and deep soils. A model version in which $m_l$ was set to zero showed negligible difference in the solutions.

#### SI Text 2: Variations in model structure

The consumption matrix dictates which populations consume which OM pools (Fig. S1). The organization of community structure and function is not well understood, and so we experiment with different possibilities. All solutions show the same general pattern: recalcitrance indicator $Q = 1$ delineates the unequilibrated from the equilibrated pools, with the latter matching the minimum subsistence concentrations among the populations. We experiment with variations in 'specialist' vs. 'generalist' capability of the populations with respect to the number of OM pools taken up by each population ($n_{up}$, which can vary from one to $n$, the number of OM pools), as well as with the widespread popularity of each pool with respect to the number of consumers of each ($n_{cons}$, which can vary from one to $m$, the number of populations) (Fig. S4).

**Default model versions (main text)** Fig. S2 shows detailed results of the two ensembles illustrated in the main text in Fig. S1. In the version with only 'specialists,' $Q < 1$ for most of the functionally recalcitrant pools because the populations associated with those pools cannot sustain themselves in the environment. With no ability to utilize other pools, these populations slowly die out over time. Because the specialist populations consuming recalcitrant pools are not fueled by biomass synthesis on other pools, the overall consumption rate of the recalcitrant pools is significantly lower (Fig. S13). Because this specialist-only configuration is more computationally efficient, we use this version to demonstrate the consistency of the dynamics when 10,000 pools of OM are resolved compared to the 10-member ensemble of 1,000 OM pools each (Fig. S3).

For the ‘generalist populations’ (Fig. S2 bottom), we vary $n_{up}$ evenly across the 1,000 additional populations, and also widely vary the probability of $n_{cons}$ (Methods). The resulting biomass concentrations are plotted as a function of the number of pools taken up in Fig. S9. About 75% of the populations retain a carbon biomass equivalent to one cell per mL after ten years (assuming 1 fmol C per cell), with the remaining 25% slowly dying out and/or rebounding as other populations die out over time.

In Figs. S3–S6, we compare these results to other possibilities as follows.

**Variation in community consumption configuration** We experiment with different ways of varying $n_{up}$ and $n_{cons}$ (Fig. S4): 1) we illustrate one of the ten simulations with generalists from the default version, 2) instead of evenly varying $n_{up}$ over the populations, we evenly vary $n_{cons}$ over the OM pools and assign a weight for $n_{up}$ and sample over each row instead of over each column. 3) We construct the matrices as in 1 and 2 but using logarithmic instead of linear ranges for the assigned distributions. 4) We correlate $n_{up}$ and $n_{cons}$ by filling in the triangle of values above or below the diagonal of the matrix. All give nearly indistinguishable results (Fig. S4), although the logarithmic ranges allow for more quasi-specialists and thus a few pools whose maximum $Q$ is below zero.

**Ratio of populations to pools** We also vary the ratio of the number of consumers to the number of OM pools (Fig. S5). As the ratio decreases, from 2:1 or 1:1 in the default versions to 1:10 to 1:100 to 1:1,000 (one generalist population), the equilibrium concentrations increase. These populations all have the same penalty for the degree of generality. (In contrast, see Fig. S6 for the results with one generalist population without this penalty.) This demonstrates that varying the ratio of populations to pools does not qualitatively change the results.

**Generalist penalty** We assign each population a penalty that increases proportionally with the number of OM pools it can consume. In reality, this penalty is not well understood, and likely differs with the particular combinations of substrate, and so we experiment with different values to understand the impact of the penalty on the model structure and solutions (Fig. S6). When the penalty for taking up more pools scales in direct proportion to the benefit (i.e. when the penalty for taking up $n$ pools is a decrease in $\rho^{\max}$ of $1/n$) both generalists and specialists are sustained, with the specifics of the competitive outcome determined by the combinations of the other parameters. This $1/n$ penalty is used in the version illustrated in the main text. When the cost of taking up more pools outweighs the benefit (a stronger penalty for generalist ability; $1/n^2$ in Fig. S6), most of the generalists are outcompeted over time by the specialists. Inversely, when the benefit of taking up more pools outweighs the cost (a weaker penalty for generalist ability; $1/\sqrt{n}$ in Fig. S6), most of the specialists are outcompeted over time. The pattern of accumulation remains robust throughout these experiments, although the weaker penalty means that the generalists are able to deplete more of the pools that would otherwise have been recalcitrant (i.e., there are more occurrences of $Q > 1$ for the same distribution of the other parameters) due to the higher frequency of faster consumption rates.

**Switching mode** In Fig. S7, we compare results to a very different model in which the microbial populations can switch back and forth between consumption of different pools. We use the same 'default' community consumption matrix for this version. Rather than simultaneously consuming multiple pools, each population consumes only the one pool that maximizes its biomass synthesis rate ($y\rho^{\max}$) at each timestep. The switching mode model is much more computationally expensive, and so we compare results after only one year of integration. The results are qualitatively similar to the version with only pure specialists (Figs. S2 and S7), because the switching populations opt against the uptake of functionally recalcitrant pools if functionally labile pools are sufficiently available.

**Transformation among pools** Using a model similar to the reduced-complexity model here, ref.[8] explored the impact of the transformation of one pool to another as an excreted waste metabolic product, rather than its full oxidation to $CO_2$. This transformation has a quantitative but not qualitative effect on the solutions, unless there is a tendency of transformation towards faster consumption rates, which we think would violate the laws of thermodynamics.

#### SI Text 3: Sensitivity to parameter values

The recalcitrance indicator (Eqn. 3 in the main text) suggests that whether or not an organic pool will accumulate in the environment due to recalcitrance depends on multiple chemical, physiological, and ecological factors (Fig. S10). The value of $Q$ increases – and thus the likelihood of recalcitrance decreases – with a faster maximum consumption rate ($\rho^{\max}$), a higher yield ($y$), a higher supply rate ($s$), and a lower loss rate ($L$). $\rho^{\max}$ varies due to varying enzymatic rates and cellular enzymatic allocation, while $y$ varies with the energetic content of a substrate and cellular maintenance requirements. The supply rate $s$ is set by the production and mortality rates of the community as well as by transport and mixing. At steady state, $L$ equals the population turnover rate, which is set by predation and viral lysis rates as well as physical transport and mixing[9]. Fig. S10 illustrates this sensitivity.

The supply rate of each OM pool moderates $Q$ indirectly via the population loss rates. With higher supply of pool $i$, the amount of biomass sustained and the specific loss rate both increase for population $j$. This simply represents the well-known phenomenon that environments with high rates of primary production sustain higher biomasses. Because larger $L_j$ can decrease $Q_i$, higher supply rates increase the likelihood of functional recalcitrance.

**Sensitivity to temperature** Temperature indirectly affects $Q$ and thus concentrations of organic matter due to its impact on metabolic rates. Specifically, temperature may impact OM dynamics in at least three ways: (1) increasing the maximum consumption rate $\rho^{\max}$, and thus increasing the maximum growth rate (2) affecting the rate of supply $s$, since in reality this supply originates from living organisms, and (3) increasing the mortality constants $m$, particularly if the activity of predators and viruses increases with temperature. In real ecosystems, (2) and (3) may be correlated. The way in which temperature will impact these parameters is unclear. In particular, it is unclear how uptake affinity may change with $\rho^{\max}$. If uptake affinity $\rho^{\max}k^{-1}$ remains constant, $k$ must also increase proportionally. Following the methodology of ref.[10], we hypothesize the potential impacts of an increase in $\rho^{\max}$ with temperature, keeping in mind the uncertainty from the other potential impacts.

For the pools that can equilibrate at subsistence concentrations, the impact of temperature is ambiguous. An increase in $\rho^{\max}$ alone should decrease the equilibrium concentrations of organic matter. However, if uptake affinity $\rho^{\max}k^{-1}$ remains constant and $k$ also increases proportionally, then there will be a negligible effect on concentrations.

For some of the functionally recalcitrant pools ($Q \leq 1$), an increase in $\rho^{\max}$ with temperature may result in a relatively small (though still nonlinear) decrease in organic matter concentration. Specifically, the rate of consumption $\rho^{\max}B$ may increase quadratically with temperature. To explain, below (Eqn. S24), we show that the processing-limited biomass $B^*_{proc}$ is directly proportional to $\rho^{\max}$. If $\rho^{\max}$ itself increases linearly over small increases in temperature, then both $v$ and $B$ may increase linear, together giving a quadratic response in the consumption rate. We can express this explicitly by considering how $\rho^{\max}$ changes with temperature as $\rho^{\max} = \rho^{\max\prime}\gamma_T$, where $\gamma_T$ is the modification of the rate due to temperature. Thus, $\rho^{\max}B^*_{proc}$ can be expressed as $\rho^{\max\prime}\gamma_T B^{*\prime}_{proc}\gamma_T = (\rho^{\max\prime}B^{*\prime}_{proc})\gamma_T^2$. However, since these recalcitrant consumption rates are generally low, the total decrease in concentration will likely be low. Furthermore, $\rho^{\max}$ may not change with temperature for some pools, such as those with organic compounds physically protected by mineral structures.

However, for other functionally recalcitrant pools for which recalcitrance is near the threshold $Q = 1$, small changes in temperature may induce a large response. Even small changes in rates can be enough to change the value of $Q$ for a pool from less than one (or approximately equal to one) to greater than one. We show, for a subset of recalcitrant pools, that warming may result in a 'tipping point:' a dramatic decrease in concentration when the threshold $Q = 1$ is crossed (Fig. S10c). This suggests the potential for the standing stock of organic matter in the oceans and soils to decrease nonlinearly with warming, with much of the newly consumed organic carbon respired and released as $CO_2$. Of course, the degree of this impact depends also on how temperature affects other ecosystem processes that set the population loss rates and other factors in $Q$.

### SI Text 4: Steady state balances

**Subsistence concentrations** The subsistence concentration $C^*_{ij}$ for OM pool $i$ and population $j$ is derived as follows. We derive it for a generalist that can consume multiple pools, and show that it reduces to the specialist's subsistence concentration (Eqn. 2). For conciseness and clarity, we neglect the probability of presence $P_j$ of the populations in the following derivation, and then add it to the final expressions. For population $j = 1$ consuming multiple pools,

$$\frac{\partial B_1}{\partial t} = \sum_{i=1}^{n_{up}} y_{i1}\rho_{i1}B_1 - L_1 B_1 \ , \tag{S1}$$

for the uptake of $n_{up}$ OM pools by population 1, where for clarity, $L_1 = m_{q_1}B_1 - m_{l_1}$ (Eqn. 6). Any pool consumed by $B_1$ may also be consumed by other populations. For pool $i = A$,

$$\frac{\partial C_A}{\partial t} = s_A - \rho^{\max}_{A1}\frac{C_A}{C_A + k_{A1}}B_1 - \underbrace{\sum_{j=2}^{n_{cons}} \rho_{Aj}B_j}_{\text{uptake by pops other than } B_1} \tag{S2}$$

for the time-averaged supply rate $s_A$ and uptake of pool $A$ by $n_{cons}$ populations. Assuming a steady state and rearranging gives the subsistence concentration $C^*_{A1}$ of pool $A$ by microbial population 1 as

$$C^*_{A1} = k_{A1}\left(\frac{\rho^{\max}_{A1}B_1}{s_A - \sum_{j=2}^{n_{cons}} \rho_{Aj}B_j} - 1\right)^{-1} \tag{S3}$$

Now we can define $q_{ij}$ as:

$$q_{A1} = \frac{\rho^{\max}_{A1}B_1}{s_A - \sum_{j=2}^{n_{cons}} \rho_{Aj}B_j} \tag{S4}$$

Thus $q_{ij}$ is a population- and OM pool-specific value. (Below (Eqn. S19), we explain how $Q_i$ is the maximum across the values of $q_{ij}$ for all populations consuming pool $i$.) Substituting $q_{A1}$ into the expression gives the subsistence concentration as:

$$C^*_{A1} = \frac{k_{A1}}{q_{A1} - 1} \tag{S5}$$

Next we derive more general expressions for $q_{ij}$ and $C^*_{ij}$ that also do not depend on the biomass concentration. Again neglecting $P_j$ in Eqn. S1 for the concentration of pool $i$, and again pulling out the uptake of population 1 from the summation gives

$$\frac{\partial C_i}{\partial t} = s_i - \rho_{i1} B_1 - \sum_{j=2}^{n_{cons}} \rho_{ij} B_j \tag{S6}$$

for uptake of pool $i$ by $n_{cons}$ populations. Assuming steady state, and multiplying through by the yield $y_{i1}$ equates the production of biomass of population 1 from pool $i$ to the supply and the uptake by other consumers as

$$y_{i1} \rho_{i1} B_1 = y_{i1}(s_i - \sum_{j=2}^{n_{cons}} \rho_{ij} B_j) \ . \tag{S7}$$

We can now substitute the above expression into Eqn. S1:

$$\frac{\partial B_1}{\partial t} = \sum_{i=1}^{n_{up}} y_{i1}(s_i - \sum_{j=2}^{n_{cons}} \rho_{ij} B_j) - L_1 B_1 \tag{S8}$$

where $n_{up}$ is the number of OM pools consumed by population 1. This can be arranged assuming steady state to give the steady state biomass concentration $B^*_1$ as

$$B^*_1 = \frac{\sum_{i=1}^{n_{up}} y_{i1}(s_i - \sum_{j=2}^{n_{cons}} \rho_{ij} B_j)}{L_1} \ . \tag{S9}$$

Substituting this into Eqn. S4 for $q_{A1}$ gives

$$q_{A1} = \frac{\rho^{\max}_{A1}}{L_1} \frac{\sum_{i=1}^{n_{up}} y_{i1}(s_i - \sum_{j=2}^{n_{cons}} \rho_{ij} B_j)}{s_A - \sum_{j=2}^{n_{cons}} \rho_{Aj} B_j} \tag{S10}$$

The production of biomass of population 1 on pool $A$ is

$$y_{A1}\rho_{A1}B_1 = y_{A1}(s_A - \sum_{j=2}^{n_{cons}} \rho_{Aj}B_j)\ . \tag{S11}$$

We can substitute this into the denominator of Eqn. S10, and also substitute in the growth balance of Eqn. S7 into the numerator of Eqn. S10 to give

$$q_{A1} = \frac{\rho_{A1}^{\max}}{L_1}\frac{\sum_{i=1}^{n_{up}} y_{i1}\rho_{i1}B_1}{\rho_{A1}B_1} \tag{S12}$$

$$= \frac{\rho_{A1}^{\max}}{L_1}\sum_{i=1}^{n_{up}} y_{i1}\frac{\rho_{i1}B_1}{\rho_{A1}B_1} \tag{S13}$$

$$= \frac{\rho_{A1}^{\max}}{L_1}\sum_{i=1}^{n_{up}} y_{i1}\frac{\rho_{i1}}{\rho_{A1}} \tag{S14}$$

We now pull the ratio of uptake for pool $i = A$ out of the summation, and sum over the remaining pools $n_{up} - 1$, where now $k$ is any other pool taken up by population 1. This gives

$$q_{A1} = \frac{\rho_{A1}^{\max}}{L_1}\left(y_{A1}\frac{\rho_{A1}}{\rho_{A1}} + \sum_{k}^{n_{up}-1} y_{k1}\frac{\rho_{k1}}{\rho_{A1}}\right) \tag{S15}$$

$$= \frac{\rho_{A1}^{\max}}{L_1}\left(y_{A1} + \sum_{k}^{n_{up}-1} y_{k1}\frac{\rho_{k1}}{\rho_{A1}}\right) \tag{S16}$$

Generalized to pool $i$ and population $j$,and adding back in the overall probability of presence $P_j$, the population- and OM pool-specific values $q_{ij}$, of which $Q_i$ is the maximum across the populations, is

$$q_{ij} = P_j\frac{\rho_{ij}^{\max}}{L_j}\left(y_{ij} + \sum_{k} y_{kj}\frac{\rho_{kj}}{\rho_{ij}}\right) \tag{S17}$$

Thus the subsistence concentration for pool $i$ and population $j$ is

$$C_{ij}^{*} = \frac{k_{ij}}{q_{ij} - 1}\ . \tag{S18}$$

Fig. S8 shows the modeled concentrations against $C^*_{ij}$. Most of the values collapse precisely along the 1:1 line. For a specialist taking up no other pools, $\sum_k y_{kj} \frac{\rho_{kj}}{\rho_{ij}} = 0$, and Eqn. S18 reduces to Eqn. 2 in the main text, and is identical to that used in a previous study for chemoautotrophic nitrifying microorganisms[11]. Thus Eqn. S18 for $C^*_{ij}$ is a general expression appropriate for diverse metabolisms and substrates. We note that the term representing the impact of another pool $k$ may also represent the impact of inorganic nutrient consumption and chemolithotrophy[12].

**Recalcitrance indicator** The expression for $Q_i$ in the main text is the maximum across many population-specific subcomponents for OM pool $i$. Here in the Supplement, we explicitly label each of these population-specific subcomponents as $q_{ij}$, for population $j$ consuming pool $i$ (Eqn. S17). The values of $q_{ij}$ were used to calculate the "community recalcitrance" in Fig. 3 in the main text. For example, if 40% of the values of $q_{ij}$ for pool $i$ are less than or equal to 1, then we can consider 40% of the populations to experience pool $i$ as recalcitrant. However, even if the "community recalcitrance" is high, it only takes one value of $q_{ij} > 1$ for the pool to be considered functionally labile overall at that time and place. Thus proximal community recalcitrance of 100% equates to functional recalcitrance. Thus the recalcitrance indicator is the maximum among the population-specific values as

$$Q_i = \max_j q_{ij} \ . \tag{S19}$$

Though this formulation is applied diagnostically in the model, it was derived with the goal of being useful for application to real systems because it requires an estimation of relative rates of uptake of multiple substrates, which may be more consistent across environments, rather than the absolute magnitude of the uptake, which depends on the local concentration.

Although $Q$ is related to fitness, investigation of competition and cooperation among populations should use $C^*_{ij}$ as the metric of fitness. Recalcitrance indicator $Q$ is always appropriate for determining whether or not a pool will accumulate. However, when multiple populations compete for the same resource, the population with the lowest subsistence concentration wins. Thus, the minimum of $C^*_{ij}$ determines the equilibrated concentration for pool $i$ (Fig. S8).

**Supply-limited subsistence concentrations** We can stringently identify the OM pools for which consumption is predominantly limited by the supply of each pool, in contrast to the handling by the cell. These supply-limited pools can be thought of as the most labile among the functionally labile pools. If the concentration $C$ of each OM pool strongly limits the rate of uptake such that $C << k$, where $k$ is the half-saturation concentration with respect to uptake, Eqn. 4 is approximated as

$$\rho_{supp_{ij}} = \rho^{\max}_{ij} \frac{C_i}{k_{ij}} \; . \tag{S20}$$

With this approximation, $q_{ij} >> 1$. This is equivalent to the approximation made in ref.[13]. With this, the population-specific supply-limited equilibrium concentration of each OM pool $C^*_{supp_{ij}}$ is:

$$C^*_{supp_{ij}} = \frac{k}{q_{ij}} \; . \tag{S21}$$

Plugging in $q$ for a specialist gives the direct comparison to Eqn. 2 in the main text:

$$C^*_{\mathrm{supp}_{ij(\mathrm{specialist})}} = \frac{k_{ij} L_j}{P_j \rho^{\max}_{ij} y_{ij}} \tag{S22}$$

Fig. S8 shows the modeled concentrations against $C^*_{supp}$. The pools closely match $C^*_{supp}$ when $Q$ is significantly greater than one. This supply-limited subset of the subsistence concentrations is analogous to the expression for total DOC found by Mentges et al. 2019[13] when summed across the pools.

**Biomass approximations** In the marine ecosystem model, we approximate the biomass associated with both the recalcitrant and supply-limited ends of the spectrum. We develop expressions for the concentrations of the biomass of the aggregated functional type populations, where each class of biomass is associated with one class of organic matter. First, we approximate the biomass sustained on recalcitrant pools – the “processing-limited” biomass – from the model parameters[8]. Because growth on the recalcitrant pools does not depend on concentration, we can make an approximation to Eqn. 6 for processing-limited biomass $B_{proc}$ as

$$\frac{\partial B_{proc_j}}{\partial t} = (y_{ij}\rho_{ij}^{\max} - m_{q_j} B_{proc_j} - m_{l_j}) B_{proc_j} \ , \tag{S23}$$

and thus uptake of pool $i$ does not depend on its concentration. We can then take the steady state approximation of Eqn. S23. Even though the associated recalcitrant pool $i$ is not at steady state (i.e. the steady state of Eqn. 5 does not approximate the solutions), this approximation ends up in a close match to the simulated biomass since the change in the biomass concentration is small relative to the rate of growth and loss of the population (Fig. S15f). From Eqn. S23 at steady state, the steady state processing-limited biomass $B^*_{proc}$ is

$$B^*_{proc_j} = (y_{ij}\rho_{ij}^{\max} - m_{l_j}) m_{q_j}^{-1} \ . \tag{S24}$$

Thus the biomass associated with recalcitrant pools reaches quasi-equilibrium, suggesting that natural assemblages may maintain steady biomass amidst varying amounts of recalcitrant OM. These results are somewhat consistent with previous work finding equilibrium relationships between OM consumption rates and biomass[14,15].

Similarly, we can approximate the supply-limited biomass $B^*_{supp}$. Using the above approximation for supply-limited consumption, $C << k$, and again assuming consumption of one OM pool for simplicity,

$$\frac{\partial B_{supp_j}}{\partial t} = (y_{ij}\rho_{ij}^{\max}\frac{C_i}{k_{ij}} - m_{q_j} B_{supp_j} - m_{l_j}) B_{supp_j} \ . \tag{S25}$$

At steady state,

$$B^*_{supp_j} = (\frac{y_{ij}\rho^{\max}_{ij}}{k_{ij}}C_i - m_{l_j})m^{-1}_{q_j} \ . \tag{S26}$$

Thus in contrast to $B^*_{proc}$, $B^*_{supp}$ must be related diagnostically to the concentration of the pool (rather than being predicted only from model parameters). However, this still provides a useful delineation of the controls on the functional biomass in the water column model (Fig. S15f).

#### SI Text 5: Simulation of Arietta experiments

To demonstrate the model consistency with the disparate hypotheses for OM concentrations, we used the model to simulate the experiments that served as evidence for dilution-limited remineralization[16] (Fig. S12). We conducted an experiment for three models: 1) the default model, 2) the model where the total number of consumers of each pool $n_{cons}$ is sampled from a linear distribution ("evenCMS" in Fig. S4, notated as "CMS" in Fig. S12), and 3) as in 2 but with 100 instead of 1,000 microbial populations. We used the "CMS" configuration to assure that each OM pool has at least one consumer for the model with 100 populations (and so experiment 2 was conducted to show consistency with experiment 1). For each experiment, we initiated the model by multiplying the previous resulting solutions after ten years of integration by a factor of ten, representing concentration of the OM. We preserved the combinations of parameters used for the interaction of each pool with each population, except for the mortality parameters. Mortality was reduced to simulate the inhibition of predation in the incubation experiments, and this impacted the concentrations of biomass but not the overall character of the solutions. Mortality rates were set to zero in the illustrated simulation. For lower mortality values, rather than zero, biomass production was slightly lower, and the evolution of the DOM concentrations remained the same. We initialized the models with 10x the concentration of the original simulated OM pools, and ran the models forward for 30 days.

Results show a continuum of fast and slow productivity (Fig. S12). The consumption of the recalcitrant pools is negligible. In consistency with the dilution hypothesis, much of the DOM is remineralized quickly, spurring microbial productivity in the transient state. A testable hypothesis arising from this is that in such an experiment, the functionally labile DOM pools that were previously equilibrated return to the steady state subsistence concentrations over a spectrum of timescales: some return relatively quickly, while some return slowly. This fuels the observed microbial growth. In contrast, the functionally recalcitrant pools remain concentrated. Furthermore, of the pools that were consumed, the timescale of consumption of each individual pool was not simply predicted from any one model parameter, such as the maximum uptake rate. Rather, the initial abundance of each heterotrophic population determined its prevalence in the newly fueled biomass production, and this abundance was set by the combination of model parameters.

We also note that because the experimental set-up alters the ecological context, increasing or decreasing the loss rates experienced by each population (which we simulated by changing the mortality parameters), both $Q$ and $C^*_{Supp}$ should also increase or decrease in such experiments relative to the *in situ* conditions. For example, reducing the mortality rate in the experimental set-up increases $Q$, which means that some pools that were functionally recalcitrant could cross the threshold and become functionally labile. This could spur even more microbial production.

### SI Text 6: Marine ecosystem model detail

**State variables and equations** The ecosystem model resolves 25 pools of DOM (both DOC and DON), 25 DOM-consuming heterotrophic functional aggregate populations, one pool of POM (both POC and PON), one POM-consuming heterotrophic population, two phytoplankton populations (one large and one small), dissolved inorganic carbon (DIC), and dissolved inorganic nitrogen (DIN). In the model, total carbon and nitrogen are both conserved over the domain. The model is of a generalized form in which carbon and nitrogen of the organic pools and the biomasses are each resolved independently, modified from ref.[8] to allow for C and N decoupling but without explicitly resolving cell quotas (detail below). This decoupling allows for the preferential consumption of recalcitrant N relative to recalcitrant C, extending the bulk turnover timescale of DOC. Here, given the quantitative uncertainties in preferential remineralization, we assume constant elemental stoichiometry to most conservatively estimate the DOC turnover timescale, without the additional impact of preferential remineralization. Therefore DON and PON remain coupled to DOC and POC, and we below list a reduced (but complete) set of equations accordingly. Parameter values are listed in Table S1.

The water column vertical domain is 2,000 m with 5 m vertical resolution. Each tracer $X$ is vertically diffused throughout the water column as function of the diffusive coefficient $\kappa$ as:

$$\frac{\partial X}{\partial t} = \nabla \cdot (\boldsymbol{\kappa} \nabla X) + S_X \qquad \text{(S27)}$$

where $S_X$ are additional sources and sinks. Sources and sinks (as well as the advection of POM according to vertical velocity $w_s$) for organic pool $i$ and population $j$ with carbon-based biomass $B_j$ are as follows:

$$S_{B_j} = \mu_j B_j - L_j B_j \tag{S28}$$

$$S_{DOC_i} = \frac{\omega_i}{\sum_{\omega_i}} \left[f_{mort} L_j B_j + (\alpha - 1)\rho_{POC,j} B_j\right] - \rho_{DOC_i,j} B_j \tag{S29}$$

$$S_{POC} = (f_{mort} - 1) L_j B_j - \alpha \rho_{POC,j} B_j - \frac{\partial(w_s POC)}{\partial z} \tag{S30}$$

$$S_{DIN} = \sum_j (\rho_{DON_i,j} - \mu_j) B_{Nj} + (\rho_{PON,j} - \mu_j) B_{Nj} - \sum_j \rho_{DIN,j} B_{Nj} \tag{S31}$$

(S32)

where $\mu_j$ is the growth rate of population $j$ (detail below for phytoplankton vs. heterotrophs), $\rho_{i,j}$ is the uptake of pool $i$ by population $j$, and $L_j = m_q B_j$. Total DOC is produced as a fraction of mortality according to $f_{mort}$ and as a fraction of POC hydrolysis according to $\alpha$. Each DOC pool is produced according to the distribution in Fig. S15d with weight $\omega_i$. Due to the assumption of constant stoichiometry, nitrogen-based biomass $B_{Nj} = B_j R_{CN}$ and organic uptake rates $\rho_{DOC} = \rho_{DON} R_{CN}$ and $\rho_{POC} = \rho_{PON} R_{CN}$ (detail below).

**Quasi-equilibrium state** All pools reach an equilibrium state except for the one pool that is recalcitrant throughout the entire water column (the lightest yellow line in Fig. S15b). Because of the unequilibrated nature of the solutions, attention to the initial conditions of the model is warranted. All pools are initialized with a constant 1 $\mu$M C at all depths, and the model is integrated forward for 6,000 years, which is more than four-fold the model equilibration timescale set by the vertical diffusion and the domain height, assuring equilibration with respect to vertical transport. At this timepoint, total DOC increases over the total domain at 0.01% per year, so the solutions may be considered quasi-steady. The only pool that continues to change in time in the solution here is the one pool that remains recalcitrant throughout the water column. This pool continues to increase because the production rate exceeds the overall consumption rate over the domain. We emphasize that this pool represents a number of OM classes and molecules in aggregate. In contrast to this increase, in the ocean, recalcitrant OM appears to decrease over time since total DOC decreases along the deep ocean trajectory[17]. This is consistent with our framework because here, we do not make any conclusion about whether recalcitrant pools in the ocean should increase or decrease, but rather that they can plausibly change over time according to the balance of any production (or other source) with any consumption (or other sink)[18]. Thus we can infer from the ocean observations that the consumption of recalcitrant DOC exceeds its production on average in the deep ocean. Since three-dimensional resolution is necessary to simulate this behavior, we leave simulation of deep DOC gradients for future work.

**Distribution of DOM** Since we know little about the rules that dictate the distribution of the forms of organic compounds within organisms, or how they flow between pools due to microbial transformation, we assume a lognormal distribution for the distribution of total DOM production among the 25 pools in accordance with theory (Fig. S15d)[19,20]. The lognormal distribution conceptually accounts for all of the processes underlying remineralization as the expected average outcome over time and space. This assumption is also mathematically appropriate since we examine only the steady state model solutions. The form of the distribution does not qualitatively change the model dynamics. For example, an even distribution gives a qualitatively similar result[8]. However, the lognormal makes for more realistic quantitative relationships among the pools, i.e. the majority of the produced DOM compounds are more labile and do not accumulate in consistency with observations[21].

**Elemental ratios** The equations represent a simplified version of a previous model that resolved cellular quotas in order to mechanistically simulate the observed patterns of DOC:DON and remineralization ratios[8]. However, in order to most conservatively estimate the age of the DOC pools, in the illustrated version we assume a constant elemental carbon to nitrogen ratio for all pools using the empirically determined average ratio for marine organic matter $R_{C:N} = 6.6$[22]. Because of the fixed stoichiometry, the uptake stoichiometry ($\rho_{DOC} : \rho_{DON}$ in Eqn. S33) always matches that of the ambient DOM. In reality, preferential consumption of recalcitrant N relative to recalcitrant C may further increase the turnover timescale of some of the recalcitrant DOC pools[8].

**Heterotrophic growth and remineralization.** The microbial heterotrophic growth rate is limited by the specific uptake rate $\rho$ of organic C or N (of either dissolved or particulate form according to Eqn. 4) as a function of carbon yield $y$ as

$$\mu_{het} = \min(y\rho_C, \rho_N)\gamma_T \ , \tag{S33}$$

where $\gamma_T$ is the modification by temperature as described below. The yield (growth efficiency) is only with respect to organic carbon because we assume that the heterotrophs do not oxidize organic N, but rather release excess N as $NH_4^+$ in the same oxidation state[8]. The excretion rate of DIC and $NH_4^+$ (i.e. remineralization) is the difference of the relevant uptake rate and the growth rate ($\rho_C - \mu$ and $\rho_N - \mu$, respectively). In reality, larger phagotrophic zooplankton significantly contribute to organic matter remineralization. Versions of the model with explicit zooplankton populations consuming microbial biomass are qualitatively similar.

**Phytoplankton growth.** Phytoplankton growth is limited by maximum growth rate $\mu^{\max}$ ($d^{-1}$), the concentration of DIN, and light, and modified by temperature ($\gamma_T$). Light limitation was parameterized using an exponential form as a function of the instantaneous photosynthetic rate $\Gamma$ ($d^{-1}$) and the Chl a to carbon ratio $\theta$ (g/g)[23,24] as:

$$\mu_P = \mu^{\max}\frac{\text{DIN}}{k_N + \text{DIN}}\left(1 - \exp\left(\frac{-\Gamma\theta}{\mu^{\max}\gamma_N\gamma_T}\right)\right)\gamma_T \tag{S34}$$

$\Gamma$ was computed as a function of photosynthetically active radiation $I$, which decreases exponentially with depth from the maximum surface flux $I_{in}$ according to the attenuation coefficient for water $k_w$. Calculation of $\Gamma$ and $\theta$ follows ref.[25] and is described in detail in ref.[11]. Maximum growth rate and half-saturation constants are computed from cell size for small and large phytoplankton populations following data-based allometric relationships[3,26,27]. Effective half-saturation constants $k_N$ for DIN limitation with respect to $\mu^{\max}$ were calculated following refs.[28,25], also described in detail in ref.[11].

**Loss rates** Loss rates of all microbial populations are parameterized using the quadratic mortality functional form to implicitly represent the impact of predation by higher trophic levels and viral lysis[29]. This is the simplest parameterization for biomass loss, and it is appropriate to choose this simple form because we lack a clear understanding of how active populations of small predators consume microbial populations (i.e. which predators consume which functional types). Furthermore, the quadratic mortality parameterization prevents the complete exclusion of any one population in an environment, which best simulates losses due to viral lysis. This results in a minimum of $Q \approx 1$. The DOC profile remains qualitatively robust across all loss parameterizations. For example, alternative versions of the model with explicit zooplankton (grazing) populations give qualitatively similar results in DOC. However, the explicit grazing results in differences in the biogeography of the functional biomass in the surface mixed layer (i.e. the slower growing biomass may be excluded at the productive surface). Given the uncertainties in loss and grazing, we leave the investigation of the impact of higher trophic levels on microbial biogeography to future work.

**Temperature** All microbial metabolic rates are modified by temperature according to non-dimensional $\gamma_T$ following the Arrhenius equation[25] as

$$\gamma_T = \tau \exp(A_E(\frac{1}{T + 273.15} - \frac{1}{T_0})) \; , \tag{S35}$$

for ambient (seawater) temperature $T$, where $T_0$ is the reference temperature, $A_E$ regulates the temperature modification, and $\tau$ normalizes the maximum value. In the water column environment, we impose an exponential temperature curve fit to observed profiles in the subtropical ocean[11], which varies from 25°C at the seasurface to 2°C at depth (see Fortran code and ref.[11]). Solutions vary quantitatively but not qualitatively for different temperature profiles.

**Vertical mixing** We simulated the surface mixed layer by varying the vertical diffusion coefficient $\kappa_Z$ from a maximum $\kappa^{\max}$ at the surface to a minimum $\kappa^{\min}$ according to length scale $z_{mld}$. To smooth over numerical error due to the fixed (no flux) boundary conditions, vertical mixing was allowed to increase at the bottom of the 2,000 m domain with a 100 m length scale, simulating a bottom boundary mixed layer. In this way, the vertical mixing coefficient $\kappa_Z$ ($m^2\ s^{-1}$) is calculated at cell faces of depth $z$ as:

$$\kappa_Z = \kappa^{\max} e^{-\frac{z}{z_{mld}}} + \kappa^{\min} + \kappa^{\max} e^{-\frac{z-H}{100}} \tag{S36}$$

where H is the height of the domain.

#### SI Text 7: Considering the particle to monomer transition

Larger organic particles are often composed of long polymers. As such, microbes must use extracellular enzymes to degrade the polymers to monomers in order to uptake the OM into their cells. This degradation process is most likely not dependent on the concentration of the polymer in the environment, and so a Michaelis-Menten form may not appropriately capture the process.

We can consider a simple description of a solid particle-resolving model for particle (or polymer) $P$ and monomer $M$, neglecting physical transport, as:

$$\frac{\partial P}{\partial t} = s_i - \beta B \tag{S37}$$

$$\frac{\partial M}{\partial t} = \beta B - \rho^{\max} \frac{M}{M+k} B \tag{S38}$$

$$\frac{\partial B}{\partial t} = y\rho^{\max} \frac{M}{M+k} B - LB \tag{S39}$$

where $\beta$ is the specific rate of particle degradation to monomer. Solving for the net change in polymer and monomer gives:

$$\frac{\partial (P+M)}{\partial t} = s_i - \rho^{\max} \frac{M}{M+k} B \tag{S40}$$

We note that Eqn. S39 is identical in form to our Eqn. 1 (when neglecting probability of presence), where the relevant OM concentration is the concentration of monomer. Since we use this equation (the rate of change of the biomass of the microbial population) to derive the recalcitrance indicator, this suggests that it is the monomer that is relevant for functional recalcitrance. Therefore, the polymer-to-monomer transition may be considered as the supply rate of the monomer, and functional recalcitrance defined in terms of the characteristics of the monomer-microbe interaction.

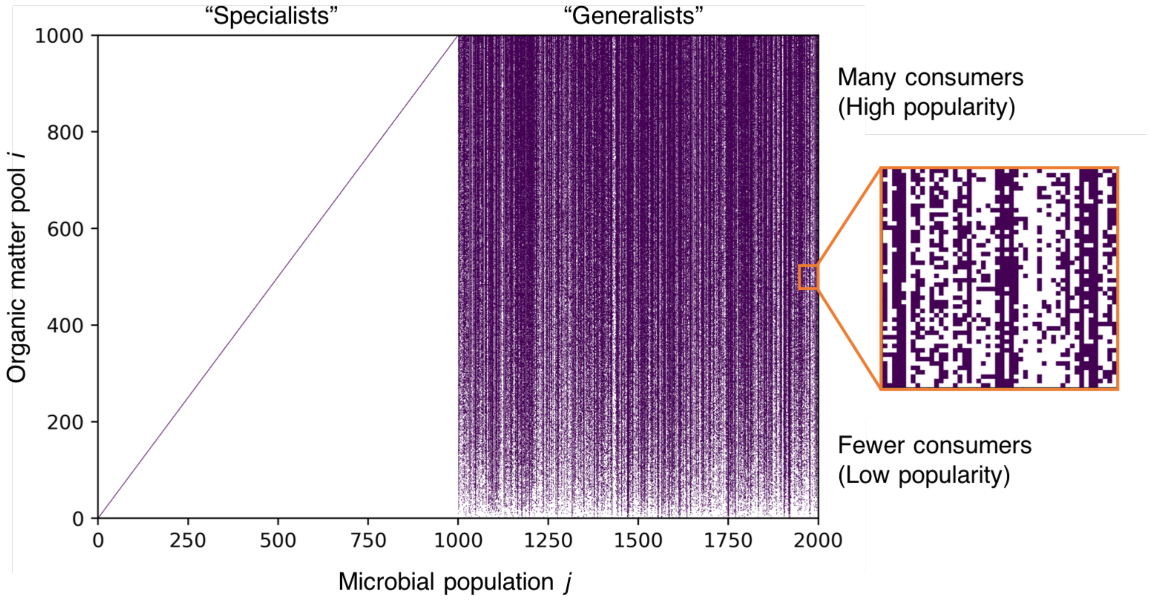


**Figure 1:** Community consumption matrix, showing the matrix for the example model version in the main text with 1,000 organic matter (OM) pools and 2,000 microbial populations. Purple colored squares indicate that population $j$ can consume OM pool $i$. The first 1,000 microbial populations are ‘specialists’ that each consume a single OM pool, which is represented by the diagonal line on the left hand side of the plot. The second 1,000 microbial populations are ‘generalists’ that are randomly assigned multiple OM pools that they are able to consume (right hand side). The number of pools consumed by each generalist $n_{up}$ varies from one to $n$, the number of OM pools. The OM pools are additionally assigned a probability of being consumed $n_{cons}$, which serves as a weight for each pool as the consumption matrix is randomly assigned (see Methods for more detail). Here, the OM pools are ordered from top to bottom according to $n_{cons}$.

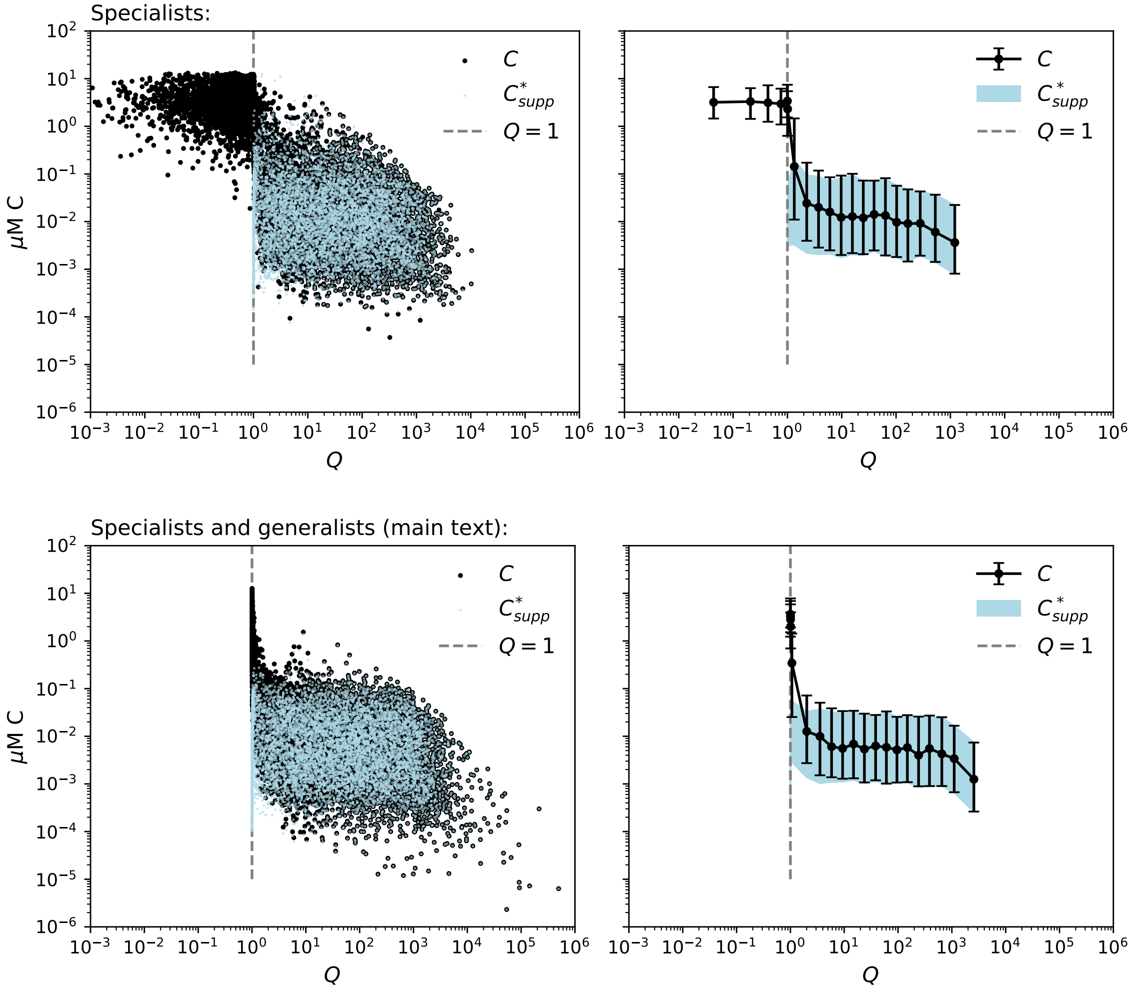


**Figure 2:** Comparison of model results with only specialist populations (top) and with both specialist and generalist microbial populations (bottom). Illustrated are the OM concentrations $C$ (black) and model diagnostic $C^*_{\rm supp}$ (the supply-limited subsistence concentrations of the microbial consumers, identifying the most labile of the functionally labile pools; Eqn. S21; light blue), against recalcitrance indicator $Q$ (Eqn. 3). $Q = 1$ (grey dashed line) separates the functionally recalcitrant OM pools from the functionally labile pools. The left plots show $C$ and $C^*_{\rm supp}$ for each individual OM pool (dots), while the right plots show the binned mean of $C$ (black line) and the 16th and 84th percentiles (equivalent to one standard deviation for a Gaussian distribution) for $C$ (black error bar) and for $C^*_{\rm supp}$ (light blue shaded region). Results from ten simulations with 1,000 OM pools each are compiled.

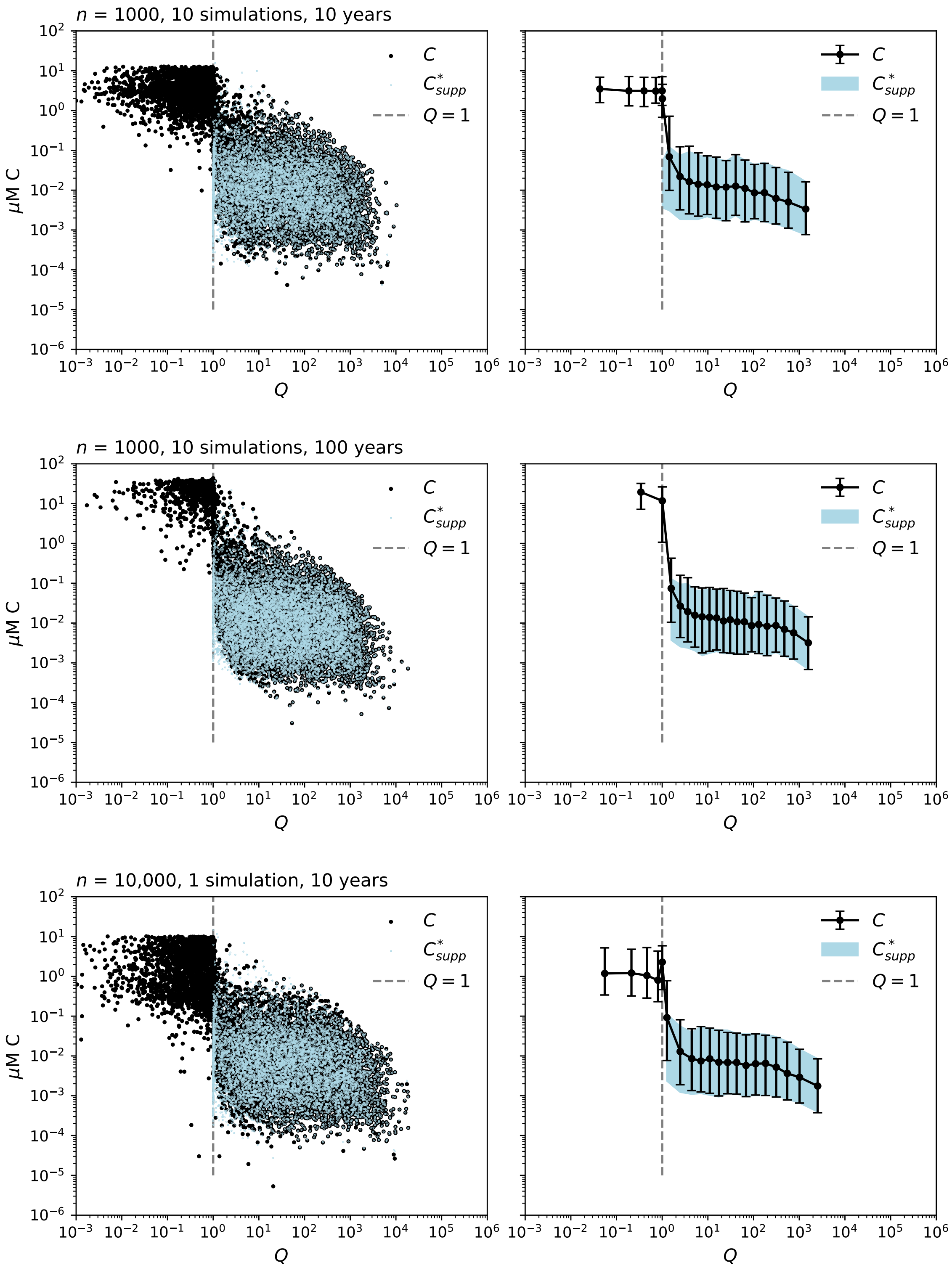


**Figure 3:** Results for specialist-only model versions, comparing the length of run time and the number of OM pools. Illustrated are the OM concentrations $C$ (black) and model diagnostic $C^*_{\mathrm{supp}}$ (the supply-limited subsistence concentrations of the microbial consumers, identifying the most labile of the functionally labile pools; Eqn. S21; light blue), against recalcitrance indicator $Q$ (Eqn. 3). $Q = 1$ (grey dashed line) separates the functionally recalcitrant OM pools from the functionally labile pools. The left plots show $C$ and $C^*_{\mathrm{supp}}$ for each individual OM pool (dots), while the right plots show the binned mean of $C$ (black line) and the 16th and 84th percentiles (equivalent to one standard deviation for a Gaussian distribution) for $C$ (black error bar) and for $C^*_{\mathrm{supp}}$ (light blue shaded region). Results from ten simulations with 1,000 OM pools each are compiled.

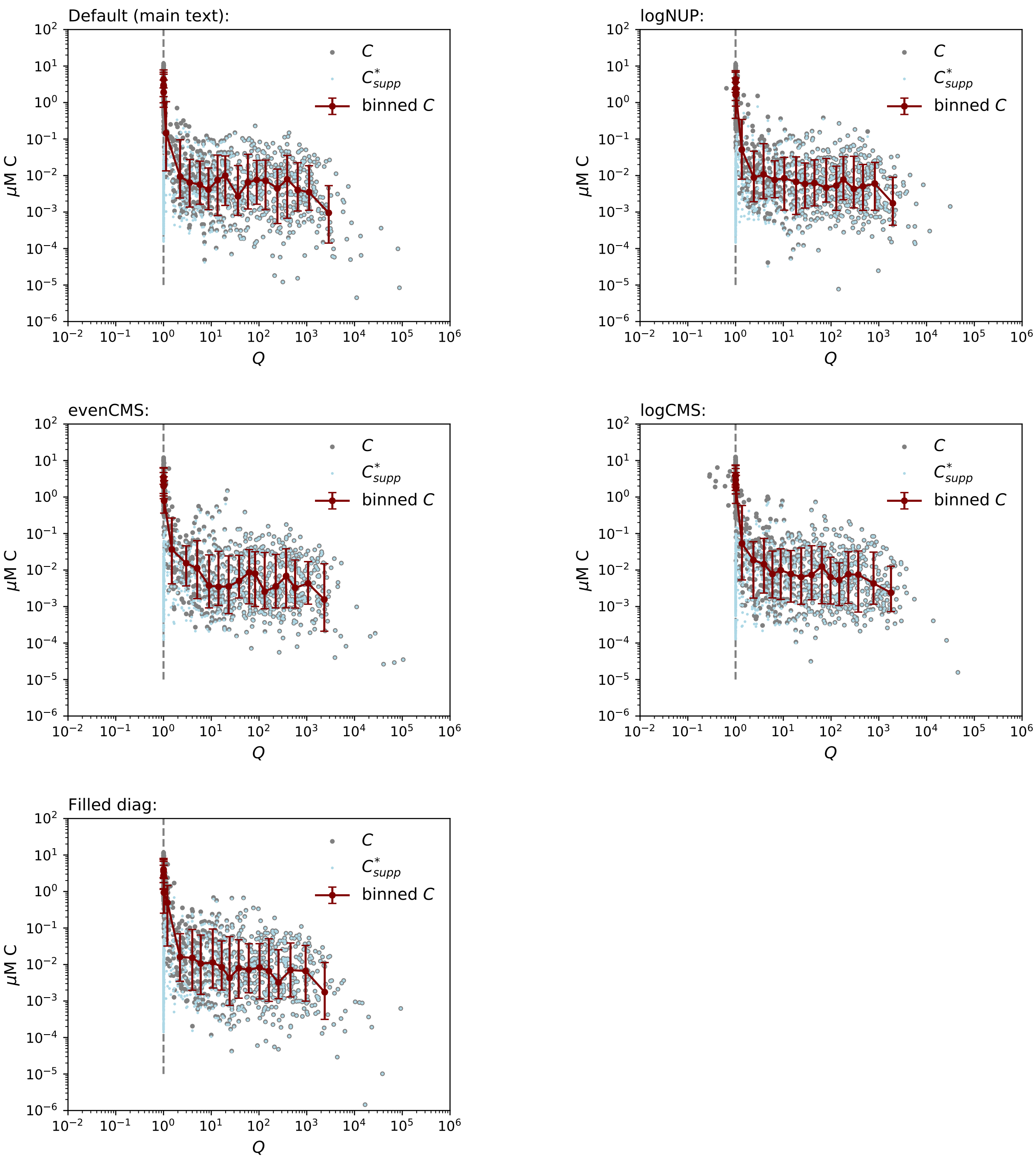


**Figure 4:** Results with different consumption configurations. *Default:* As in the main text, where the number of pools taken up by each population $n_{up}$ is sampled randomly from a linear distribution. *logNUP:* $n_{up}$ is sampled from a logarithmic distribution. *evenCMS:* The total number of consumers of each pool $n_{cons}$ is sampled from a linear distribution. *logCMS:* $n_{cons}$ is sampled from a logarithmic distribution. *Filled diag:* $n_{up}$ and $n_{cons}$ are correlated by filling the lower diagonal of the consumption matrix. Illustrated are the individual OM concentrations $C$ (grey dots), the individual model diagnostic $C^*_{\mathrm{supp}}$ (the supply-limited subsistence concentrations of the microbial consumers, identifying the most labile of the functionally labile pools; Eqn. S21; light blue dots), the binned mean of $C$ (maroon line), and the 16th and 84th percentiles (equivalent to one standard deviation for a Gaussian distribution) for $C$ (maroon error bar) against recalcitrance indicator $Q$ (Eqn. 3). $Q = 1$ (grey dashed line) separates the functionally recalcitrant OM pools from the functionally labile pools. Each simulation has 1,000 OM pools. See SI Text 2 for more detail.

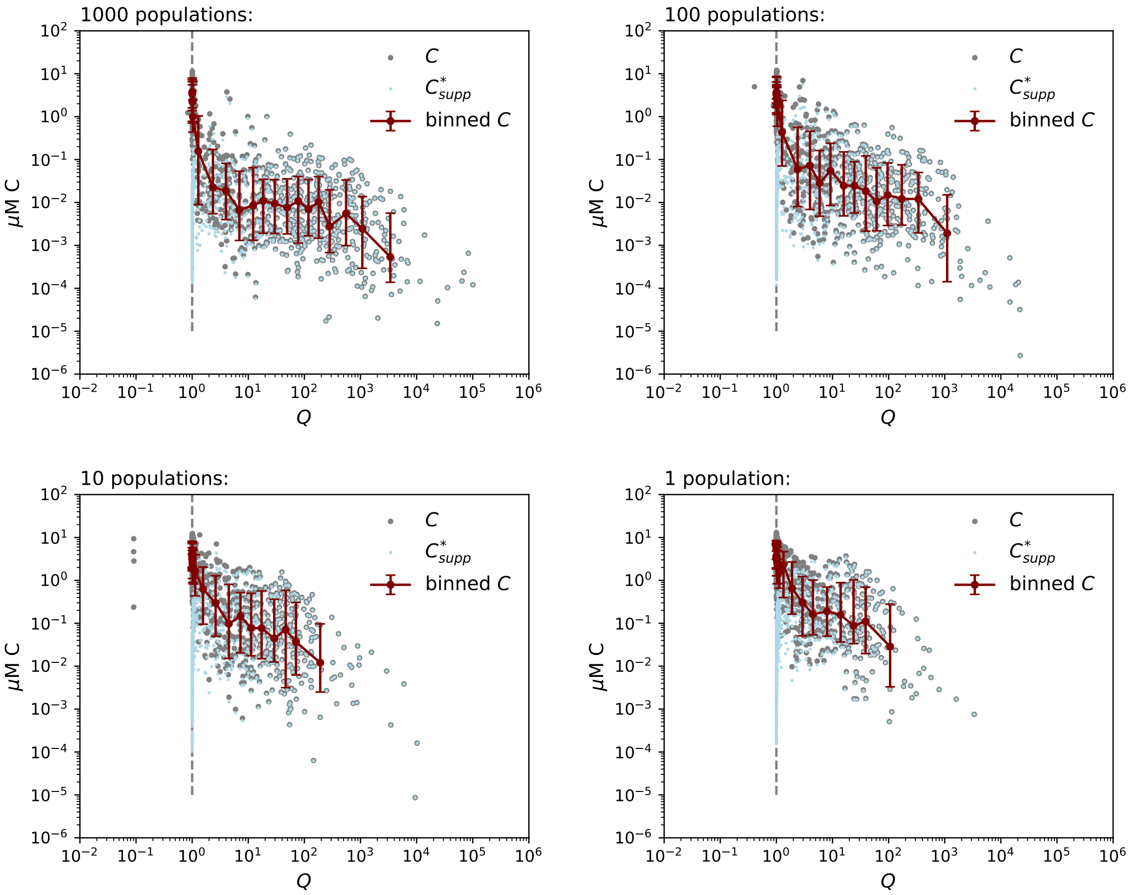


**Figure 5:** Results with variation in the number of microbial populations, and thus the ratio of populations to OM pools. Each simulation has 1,000 OM pools. Illustrated are the individual OM concentrations $C$ (grey dots), the individual model diagnostic $C^*_{\text{supp}}$ (the supply-limited subsistence concentrations of the microbial consumers, identifying the most labile of the functionally labile pools; Eqn. S21; light blue dots), the binned mean of $C$ (maroon line), and the 16th and 84th percentiles (equivalent to one standard deviation for a Gaussian distribution) for $C$ (maroon error bar) against recalcitrance indicator $Q$ (Eqn. 3). $Q = 1$ (grey dashed line) separates the functionally recalcitrant OM pools from the functionally labile pools.

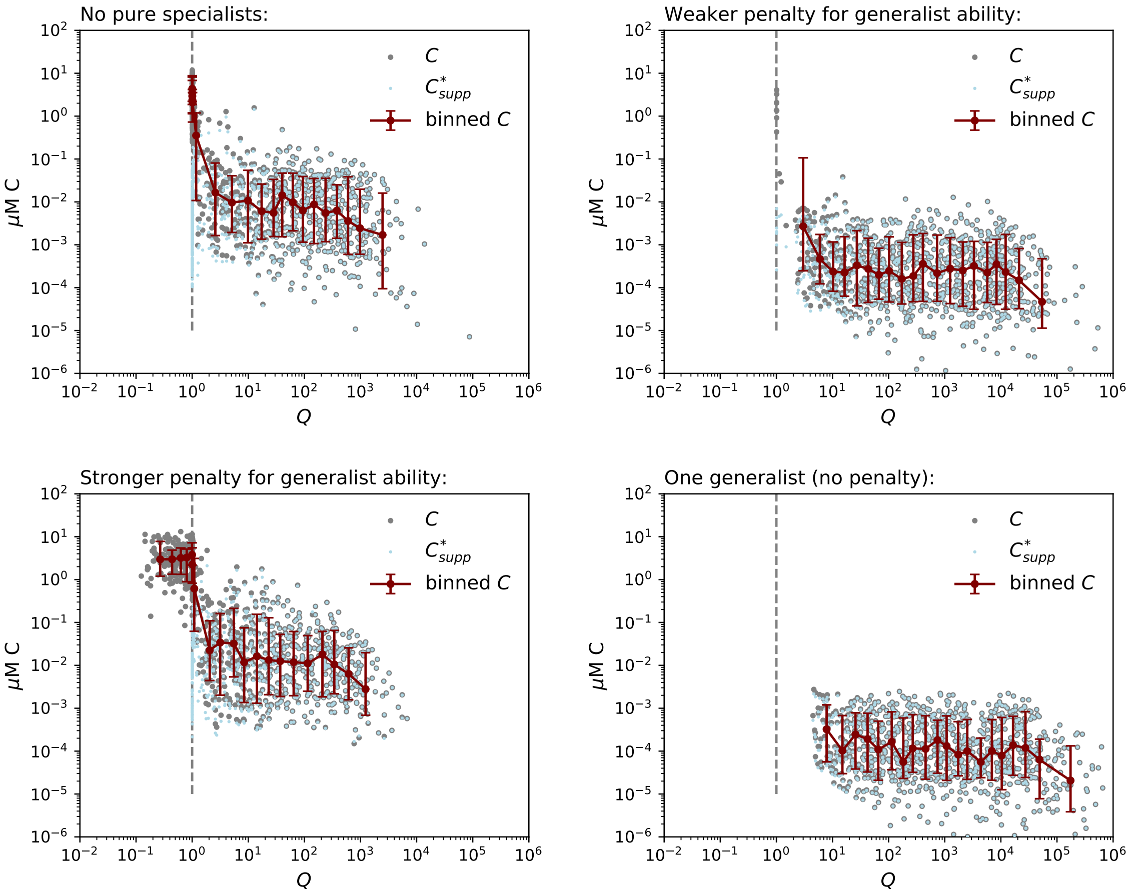


**Figure 6:** Comparison of different penalties for generalists in the model. In the model version illustrated in the main text, the penalty for taking up $n_{up}$ pools is a decrease in $\rho^{\max}$ of $1/n_{up}$. *No pure specialists*: the same penalty as in the model illustrated in the main text (a decrease in $\rho^{\max}$ of $1/n_{up}$), but here with only 1,000 generalist populations for better comparison of the different penalties. *Weaker penalty for generalist ability*: The decrease in $\rho^{\max}$ is $1/\sqrt{n}_{up}$. *Stronger penalty for generalist ability*: The decrease in $\rho^{\max}$ is $1/n_{up}^2$. *One generalist (no penalty)*: Only one population consumes all pools, with no decrease in $\rho^{\max}$. In other words, the magnitudes of the rates remain the same as for the model with pure specialists, except that they all are associated with uptake by one population of biomass. Illustrated are the individual OM concentrations $C$ (grey dots), the individual model diagnostic $C^*_{\text{supp}}$ (the supply-limited subsistence concentrations of the microbial consumers, identifying the most labile of the functionally labile pools; Eqn. S21; light blue dots), the binned mean of $C$ (maroon line), and the 16th and 84th percentiles (equivalent to one standard deviation for a Gaussian distribution) for $C$ (maroon error bar) against recalcitrance indicator $Q$ (Eqn. 3). $Q = 1$ (grey dashed line) separates the functionally recalcitrant OM pools from the functionally labile pools. Each simulation has 1,000 OM pools.

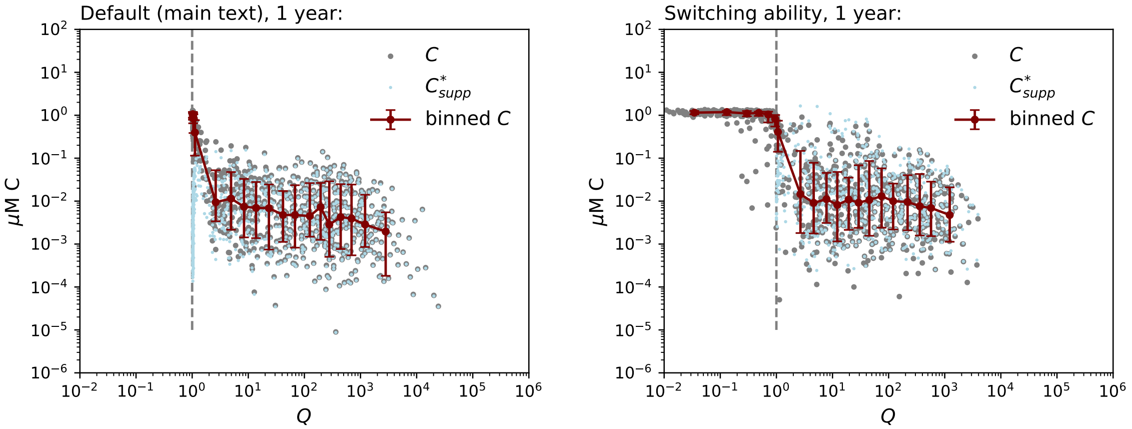


**Figure 7:** Comparison of two model versions: where the generalists are able to take up all pools continually (*Default*), and where the generalists are able to switch and take up only the pool that allows them the maximum biomass synthesis rate at each timestep (*Switching ability*). Integration time is one year because of the computational expense of selecting the optimal pool for growth. Illustrated are the individual OM concentrations $C$ (grey dots), the individual model diagnostic $C^*_{\text{supp}}$ (the supply-limited subsistence concentrations of the microbial consumers, identifying the most labile of the functionally labile pools; Eqn. S21; light blue dots), the binned mean of $C$ (maroon line), and the 16th and 84th percentiles (equivalent to one standard deviation for a Gaussian distribution) for $C$ (maroon error bar) against recalcitrance indicator $Q$ (Eqn. 3). $Q = 1$ (grey dashed line) separates the functionally recalcitrant OM pools from the functionally labile pools. Each simulation has 1,000 OM pools.

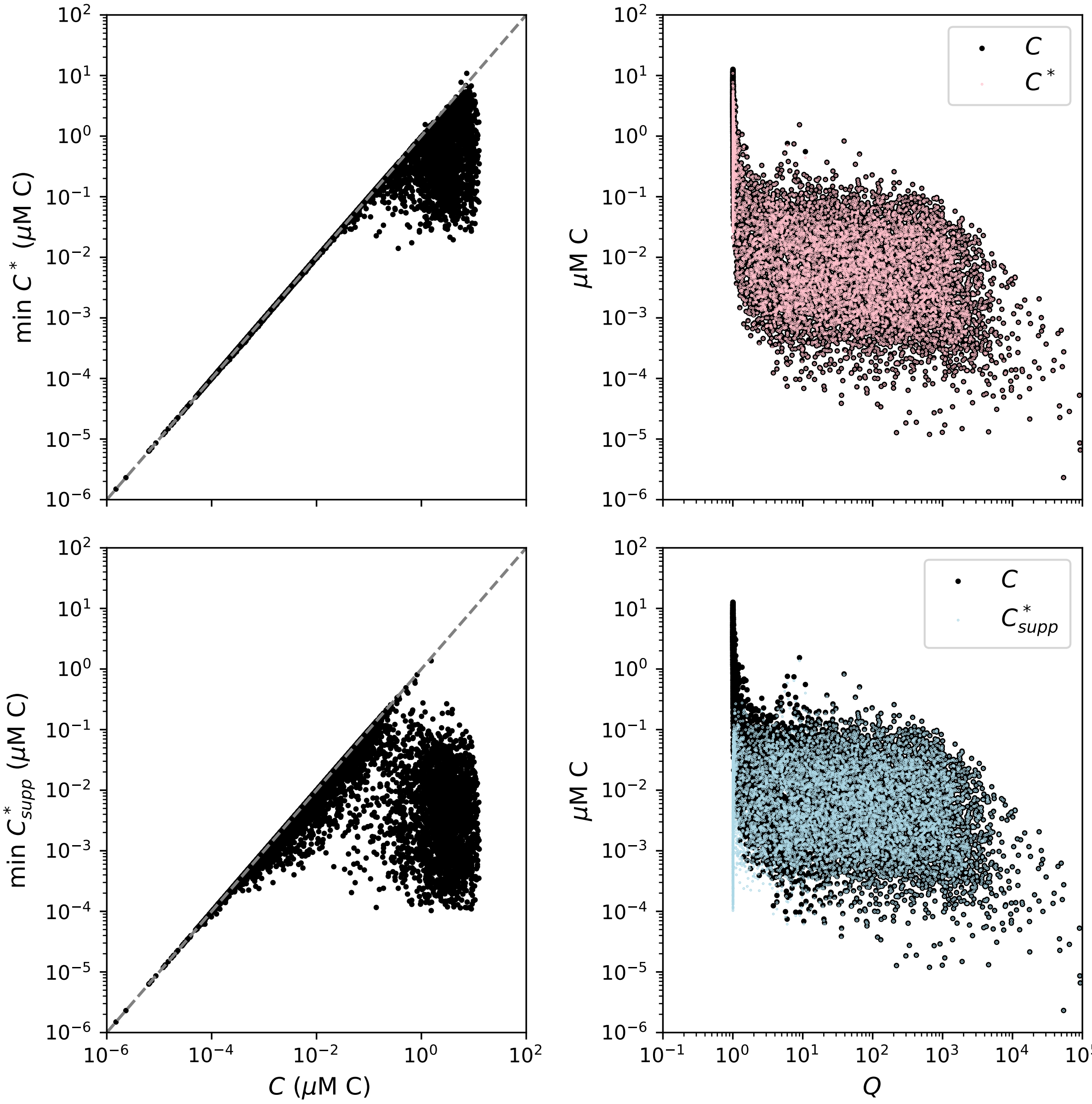


**Figure 8:** Comparison of the resulting carbon concentrations $C$ of the individual OM pools from the model solutions against the diagnostic subsistence concentrations of the microbial populations. The top row compares the resulting $C$ to the minimum of the subsistence concentrations $C^*$ (Eqn. 2, Eqn. S18) among the consumers. The bottom row compares the resulting $C$ to the minimum of the supply-limited subsistence concentrations $C^*_{supp}$ (identifying the most labile of the functionally labile pools; Eqn. S21) among the consumers. The right side plots the solutions (black dots) and the approximations (colored dots) against the recalcitrance indicator $Q$ (Eqn. 3), which delineates the functionally recalcitrant OM pools and the functionally labile pools. Results from ten simulations with 1,000 OM pools each are compiled.

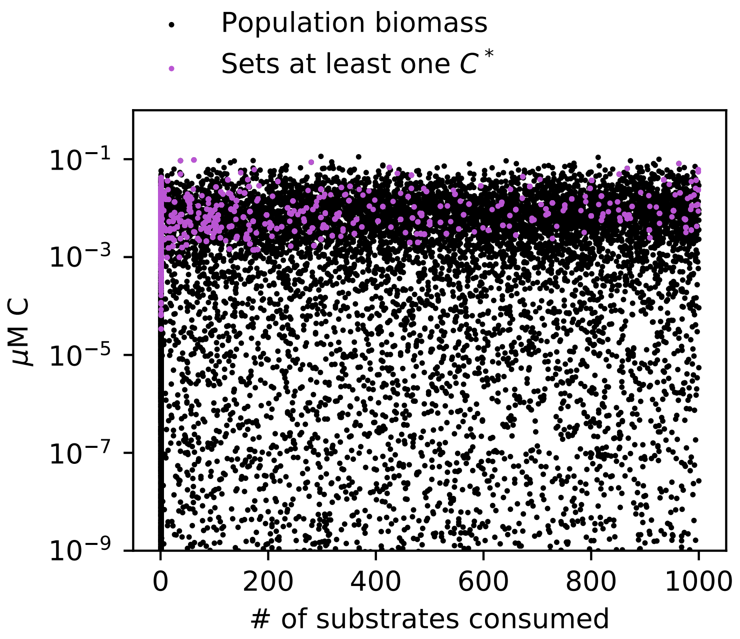


**Figure 9:** Resulting concentrations of microbial biomass in the ten year model simulation. The black dots represent the concentrations of each of the 2,000 microbial populations (1,000 specialists and 1,000 generalists). The concentrations are sorted against the number of substrates (OM pools) consumed by each population, ranging from 1 to 1,000. Thus the concentrations of specialists are all collapsed along one column (one substrate consumed). The populations whose diagnostic subsistence concentrations $C^*$ precisely match at least one of the OM pool concentrations are indicated in purple. Consumption by these populations sets the resulting equilibrated concentrations of functionally labile OM.

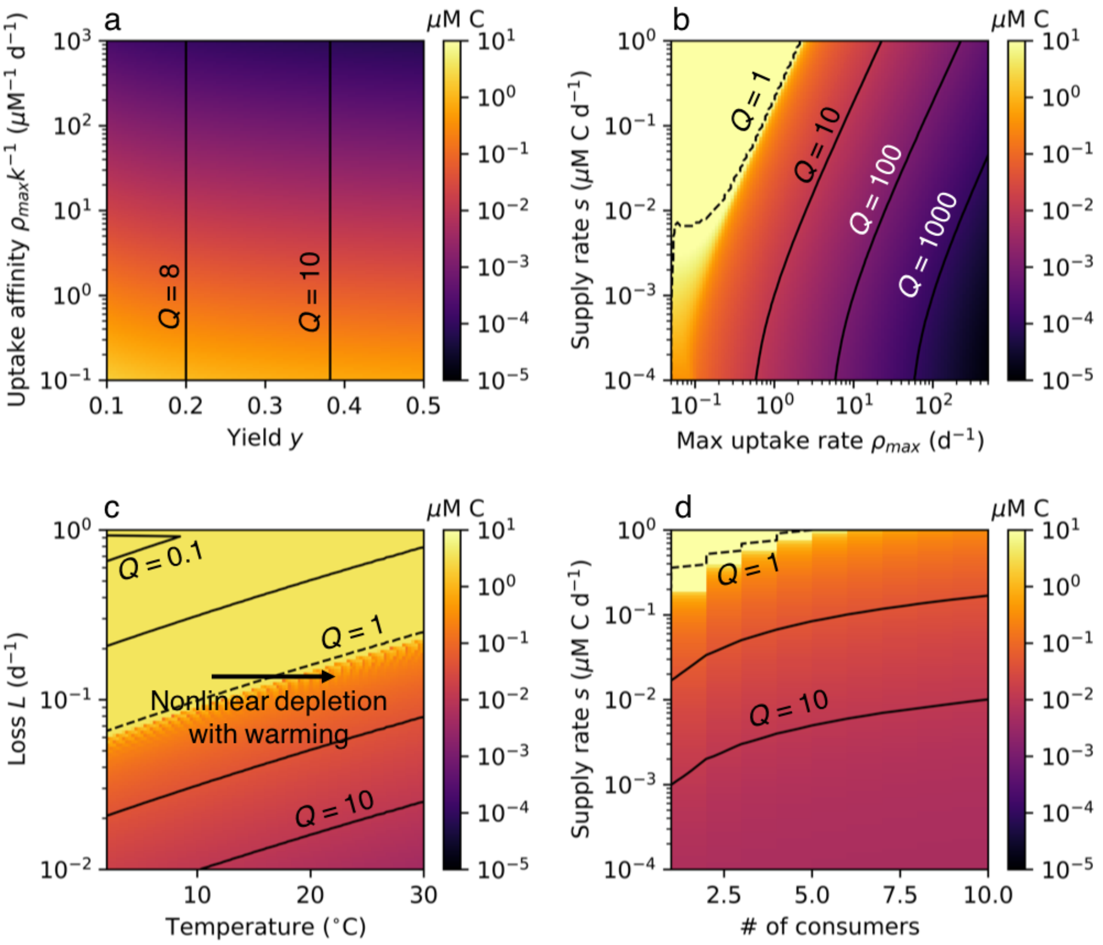


**Figure 10:** Sensitivity of organic matter (OM) carbon concentration to the model parameters. Contours (black lines) indicate values of recalcitrance indicator $Q$ (Eqn. 3). $Q = 1$ (black dashed lines) separates the accumulating, functionally recalcitrant OM pools from the equilibrated, functionally labile pools. For temperature (panel c), the maximum uptake rate increases with temperature according to the temperature modification factor $\gamma_T$ (Eqn. S34). Nonlinear responses to changes in parameter values occur when the threshold $Q = 1$ is crossed. Thus, a nonlinear depletion in carbon concentration may result for some OM pools when temperature is increased.

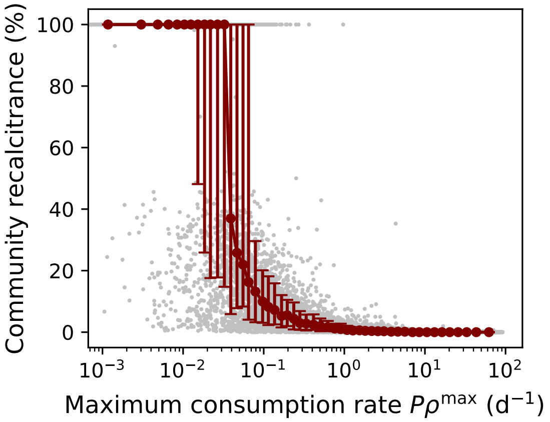


**Figure 11:** Variable 'community recalcitrance' in the stochastic microbial consumption model, defined as the percentage of consumers for which an OM pool is functionally recalcitrant. Community recalcitrance is calculated using recalcitrance indicator $Q$ (Eqn. 3) and plotted against the local maximum consumption rate of each OM pool, $P\rho^{\max}$, which is the highest combination of probability of presence $P$ and maximum uptake rate $\rho^{\max}$ among all of the populations able to consume that pool. The scatter illustrates how the multiple parameters comprising the recalcitrance indicator $Q$ contribute to functional recalcitrance. For example, for local maximum consumption rates of about 0.1 $d^{-1}$, about 5% to 20% of populations experience a particular pool as recalcitrant. This anticipates the observed slow consumption rates of substrates deemed recalcitrant. The grey dots indicate each of the 10,000 pools in the compilation of ten simulations. The larger red dots indicate the binned mean, and the red bars indicate the binned 16th and 84th percentiles (equivalent to one standard deviation for a Gaussian distribution).

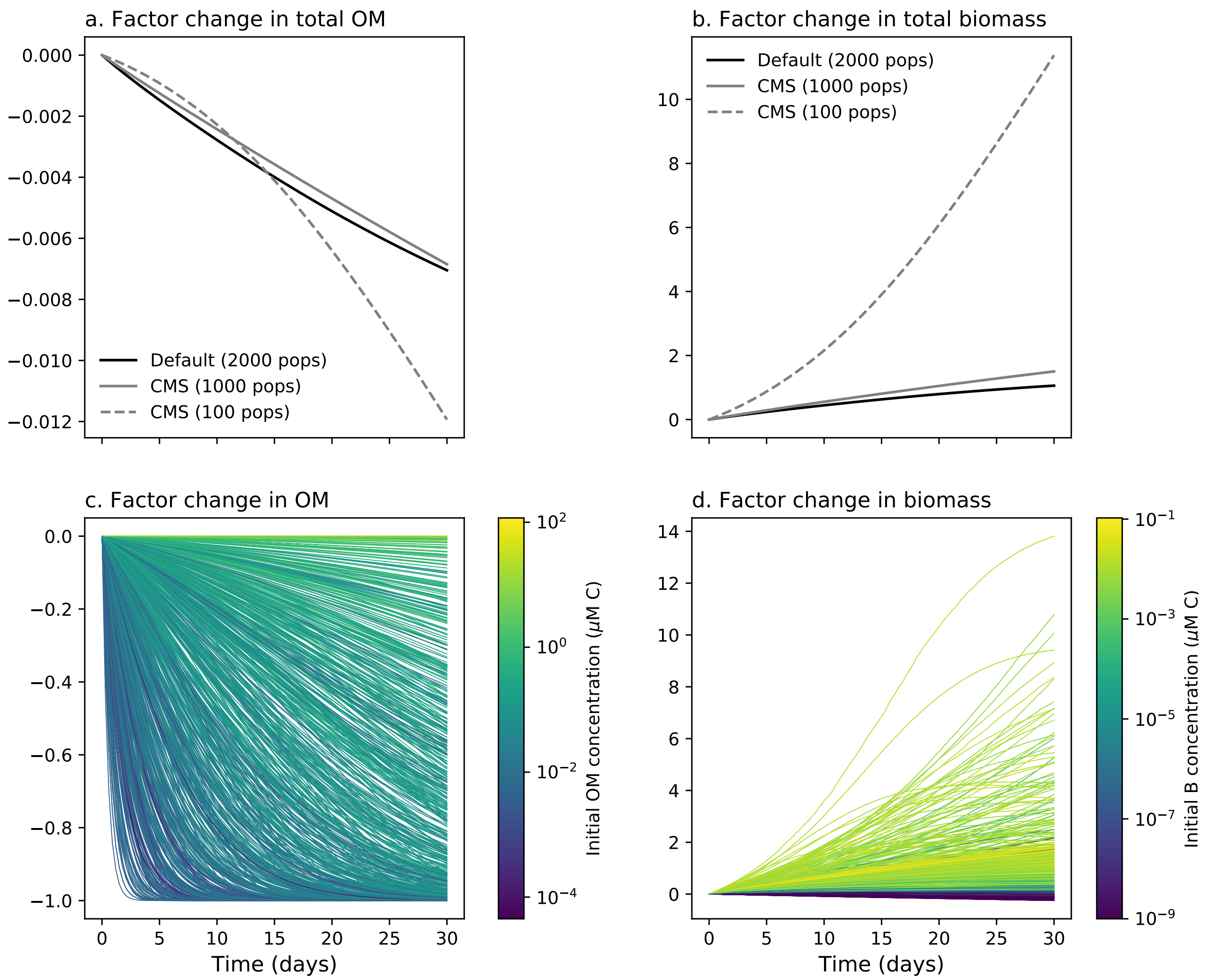


**Figure 12:** Model simulation of the experiments of Arrieta et al. 2015[16] examining the response to a ten-fold concentration of the organic matter (OM) pools. (See SI Text 5 for detail and discussion.) The default model version has 1,000 specialist and 1,000 generalist populations. To show the impact of different ratios of populations to OM pools, we show two other versions with the "CMS" configuration, with 1,000 and 100 generalist populations, because this configuration assures that each pool has at least one consumer as the number of populations is decreased (Fig. S4). As the number of populations decreases, the amount of biomass increases because each population consumes a greater number of pools. Panels a and b show the factor change in total OM and biomass for the three model versions. Panels c and d show the factor change in the individual concentrations of OM and biomass for the default model version.

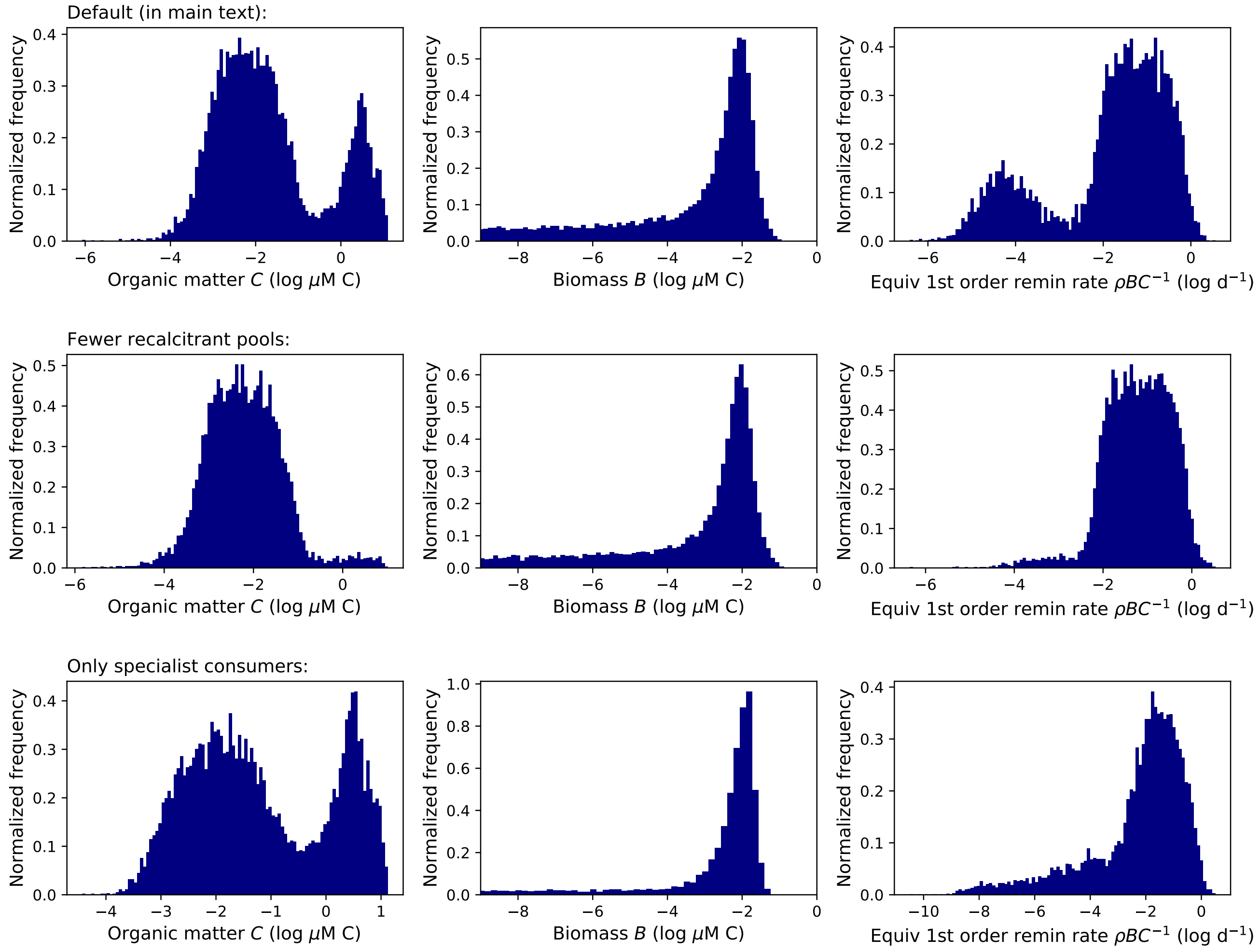


**Figure 13:** Distributions of organic matter (OM) concentrations, biomass concentrations, and remineralization rates from simulations with three different model versions. *Default (in main text):* Maximum uptake rate $\rho^{\max}$ is sampled from a distribution of $10^{-2}$ to $10^{2}$ d$^{-1}$ (Table 1). *Fewer recalcitrant pools*: $\rho^{\max}$ is sampled from a narrower distribution of $10^{-1}$ to $10^{2}$ d$^{-1}$, resulting in less functionally recalcitrant OM pools. *Only specialist consumers*: Each population is associated with the consumption of only one OM pool. Thus consumers of recalcitrant pools are not fueled by other non-recalcitrant pools, and so consumption of many recalcitrant pools ceases because more populations become extinct.

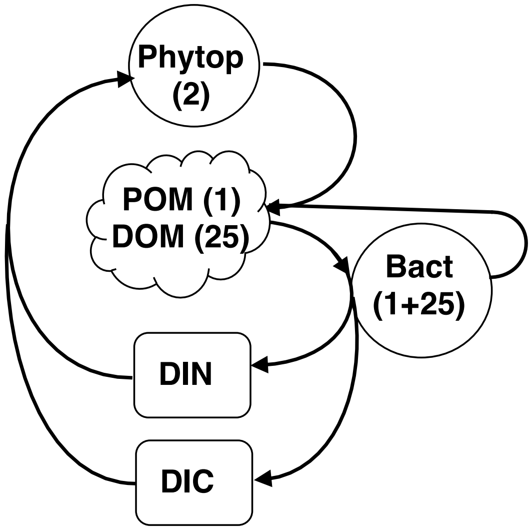


**Figure 14:** Schematic of the food web in the marine ecosystem model. The model resolves two clades of phytoplankton (one fast, one slow), one bulk pool of POM (particulate organic matter) consumed by one particle-associated bacterial clade, 25 pools of DOM (dissolved organic matter) each consumed by a unique bacterial clade, DIN (dissolved inorganic nitrogen), and DIC (dissolved inorganic carbon). Both carbon and nitrogen are resolved for the organic pools (i.e. POC, PON, DOC, and DON).

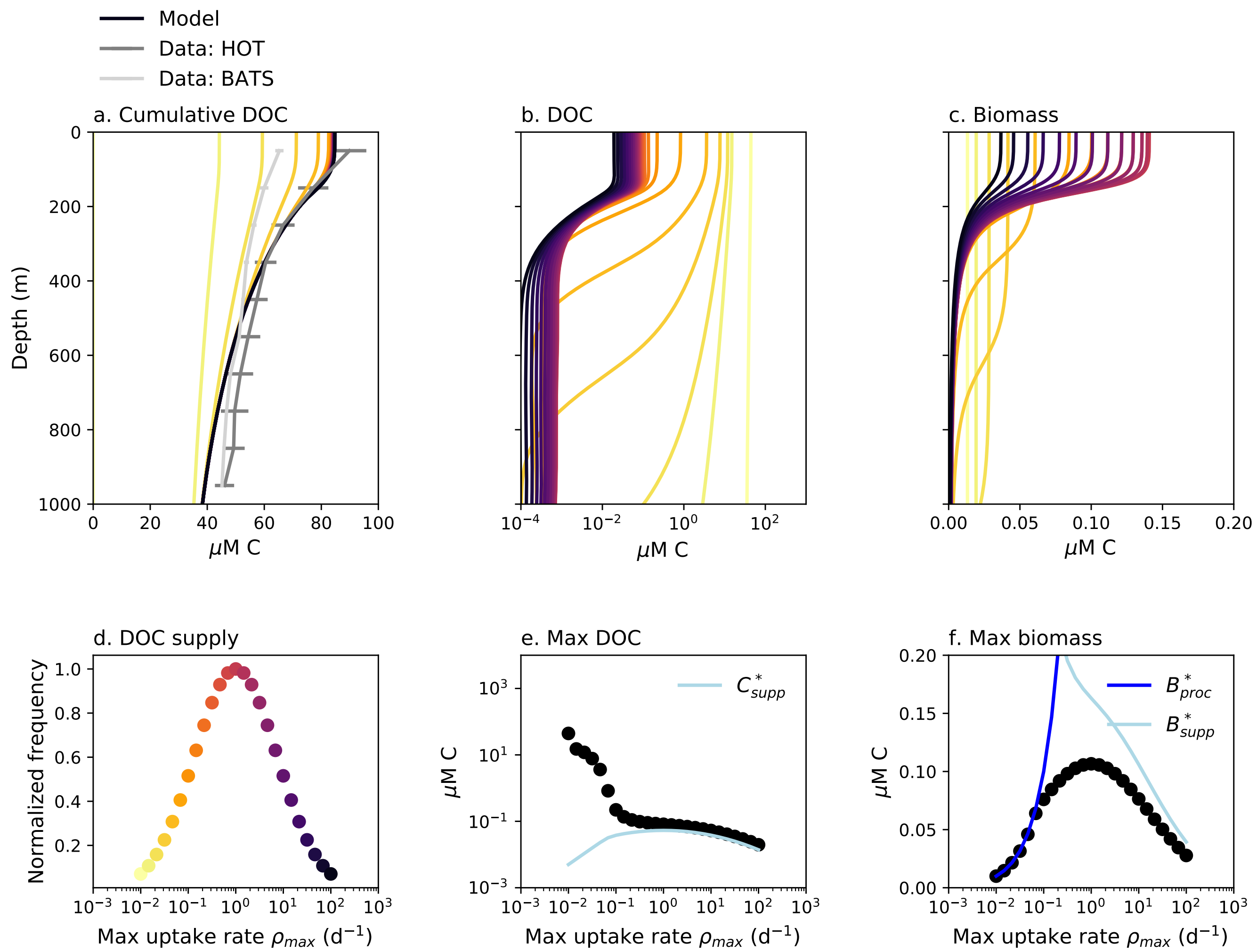


**Figure 15:** Extended water column model solutions. (a). Total modeled (black line) and observed total DOC (grey lines) concentration. Additionally, the concentrations of the 25 modeled DOC pools are plotted cumulatively (colored lines), so that the contribution to total DOC by each pool is visible, with the color in panel d serving as the legend. Observations shown are 8-year annual averages from two ocean time series: HOT (the Hawaii Ocean Time-series in the Pacific Ocean) and BATS (the Bermuda Atlantic Time-series Station in the Atlantic Ocean)[30]. (b) Concentrations of the 25 individual DOC pools. (c) Concentrations of the 25 biomass pools, with color indicating the OM pool each consumes. (d) The frequency of supply of each of the 25 pools as a function of maximum uptake rate $\rho_{max}$. The DOM is sourced from the mortality of all populations and the hydrolysis of POM (particulate organic matter). (e) The maximum (surface) concentration of each of the 25 DOC pools (black dots) and the model diagnostic $C^*_{\mathrm{supp}}$ (the supply-limited subsistence concentrations of the microbial consumers, identifying the most labile of the functionally labile pools; Eqn. S21; light blue line). (f) The maximum biomass concentration associated with consumption of each DOC pool (black dots) compared to model diagnostics: the processing-limited biomass concentration $B^*_{\mathrm{proc}}$ (Eqn. S24; darker blue line) and the supply-limited biomass concentration $B^*_{\mathrm{supp}}$ (Eqn. S26; lighter blue line). Though the recalcitrant pools are not equilibrated, the biomass clades consuming the recalcitrant pools do reach quasi-equilibrium ($B_{proc}$)[8], suggesting that natural assemblages may maintain steady biomass amidst varying amounts of recalcitrant OM. The color in panel d serves as the legend for panels a–c.

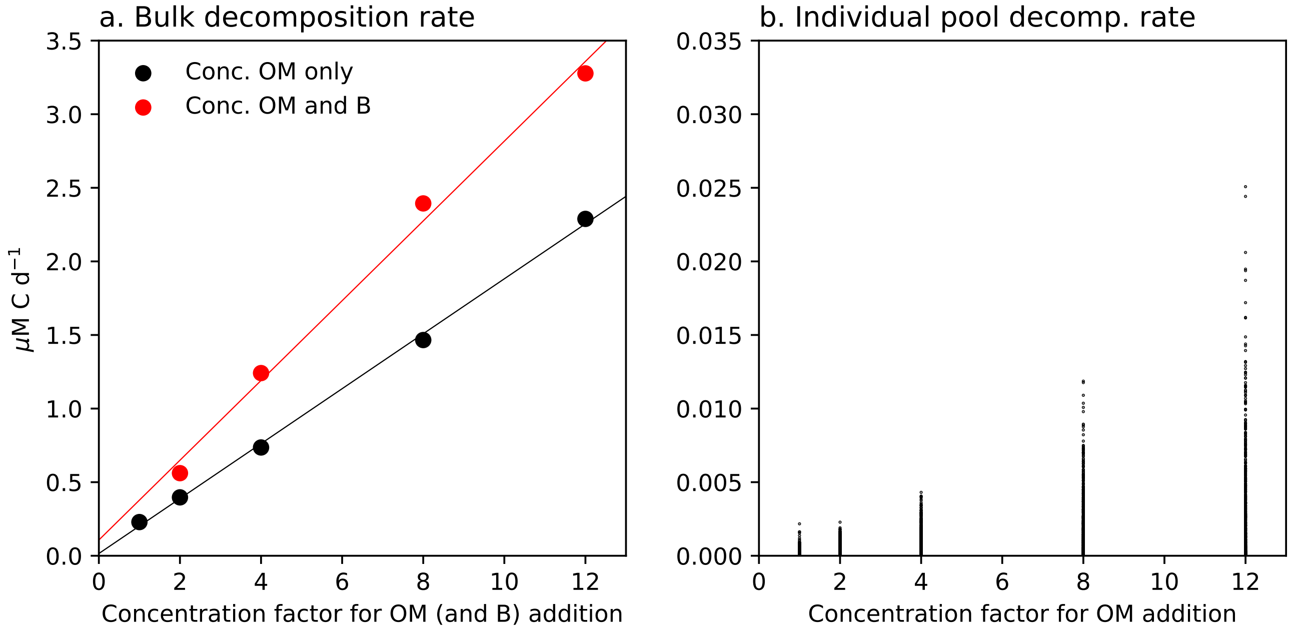


**Figure 16:** Simulation of the experiment measuring bulk sediment decomposition rate after additions of organic matter[31]. In their experiment, Westrich and Berner (1984) observed that the rate of organic carbon decomposition increased linearly with the amount of organic carbon added after ten days (their Fig. 4). From this, they confirmed the utility of first-order decomposition models (or, "multi-G" models) for describing the decomposition. Here, we show that our model, with nonlinear (Michaelis-Menten) uptake by microbial populations, also results in this bulk linear behavior. We replicated the experiment by adding different amounts of organic matter (OM) to the resulting solutions from the default model ($n$ = 1,000), and integrating the model for ten days. The additions were accomplished by initiating the model with the original solutions concentrated by a range of factors, as indicated by the x-axis. (a) The bulk rate of decomposition (calculated as the sum of the consumption by all populations of all pools; Eqn. 5) increases linearly with the additions (black dots in panel a). To test whether additions of microbial biomass impacted the results, we also conducted the experiment in which biomass concentrations were increased by the same factor as OM (red dots in panel a). Best fit linear regressions demonstrate the linearity (black and red lines in panel a, $R^2 = 0.99$ for both). (b) The decomposition rates of the individual OM pools for the experiment with only the OM additions (calculated as the sum of the consumption by all populations for each pool; Eqn. 5).

**Table 1:** Parameters for the marine water column ecosystem model.

| Parameter | Symbol | Value | Units |
|---|---|---|---|
| **Heterotrophic growth:** | | | |
| Maximum DOM uptake rate (Fig. S15d) | $\rho_i^{\max}$ | 0.01 – 100 | $d^{-1}$ |
| DOC half-saturation | $k_{\mathrm{DOC}_i}$ | $0.1\rho_i^{\max} R_{\mathrm{CN}}$ | µM C |
| Maximum POM uptake rate | $\rho_{\mathrm{POM}}^{\max}$ | 1 | $d^{-1}$ |
| POC half-saturation | $k_{\mathrm{POC}}$ | $0.1\rho_{\mathrm{POM}}^{\max} R_{\mathrm{CN}}$ | µM C |
| Carbon yield | $y$ | 0.2 | unitless |
| Production of DOM from POM hydrolysis | $\alpha$ | 15% | % of hydrolysis to DOM |
| **Phytoplankton growth:** | | | |
| Maximum growth rate, large | $\mu_j^{\max}$ | 2 | $d^{-1}$ |
| Maximum growth rate, small | $\mu_j^{\max}$ | 0.515 | $d^{-1}$ |
| DIN half-saturation, large | $k_{N_j}$ | 3.3 | µM |
| DIN half-saturation, small | $k_{N_j}$ | 3.6 | nM |
| Photosynthetic rate | $\Gamma$ | † | $d^{-1}$ |
| Chl:C | $\theta$ | † | g Chl $g^{-1}$ C |
| **Stoichiometry:** | | | |
| Biomass carbon to nitrogen ratio | $R_{\mathrm{CN}}$ | 6.6[§] | mol/mol |
| **Mortality:** | | | |
| Quadratic mortality rate | $m_q$ | 1 | µM $N^{-1}$ $d^{-1}$ |
| Fraction of mortality to DOM vs. POM | $f_{mort}$ | 0[‡] | unitless |
| **Temperature dependence:** | | | |
| Reference temperature | $T_0$ | 293.15 | K |
| Temperature regulation | $A_E$ | -4000 | K |
| Temperature normalization | $\tau$ | 0.8 | unitless |
| **Physical parameters:** | | | |
| POM sinking velocity | $w_s$ | 10 | m $d^{-1}$ |
| Surface incoming PAR flux | $I_{in}$ | 700 | W $m^{-2}$ |
| PAR attenuation in water | $k_w$ | 0.04 | $m^{-1}$ |
| Mixed-layer attenuation depth | $z_{mld}$ | 20 | m |
| Minimum vertical mixing coefficient | $\kappa^{\min}$ | $10^{-4}$ | $m^2$ $s^{-1}$ |
| Maximum vertical mixing coefficient | $\kappa^{\max}$ | $10^{-2}$ | $m^2$ $s^{-1}$ |

† Calculated as in Zakem et al. 2018[11] following Dutkiewicz et al. 2015[25].

[§] See Zakem and Levine 2019[8] for dynamics of variable C:N.

[‡] The DOC profiles are insensitive to whether sources from excretion or mortality: sensitivity analysis in Zakem and Levine 2019[8].
